# A stabilized MERS-CoV spike ferritin nanoparticle vaccine elicits robust and protective neutralizing antibody responses

**DOI:** 10.1101/2024.07.01.601243

**Authors:** Abigail E. Powell, Hannah Caruso, Soyoon Park, Jui-Lin Chen, Jessica O’Rear, Brian J. Ferrer, David M. Belnap, Audrey Walker, Anneliese Bruening, Airn Hartwig, Vida Ahyong, Cristy S. Dougherty, Richard Bowen, Julie E. Ledgerwood, Michael S. Kay, Payton A.-B. Weidenbacher, Brad A. Palanski

## Abstract

Middle East respiratory syndrome coronavirus (MERS-CoV) was first identified as a human pathogen in 2012 and causes ongoing sporadic infections and outbreak clusters. Despite case fatality rates (CFRs) of over 30% and the pandemic potential associated with betacoronaviruses, a safe and efficacious vaccine has not been developed for prevention of MERS in at-risk individuals. Here we report the design, in vitro characterization, and preclinical evaluation of MERS-CoV antigens. Our lead candidate comprises a stabilized spike ectodomain displayed on a self-assembling ferritin nanoparticle that can be produced from a high-expressing, stable cell pool. This vaccine elicits robust antibody titers in BALB/c mice as measured by MERS-CoV pseudovirus and live-virus neutralization assays. Immunization of non-human primates (NHPs) with a single dose of Alhydrogel-adjuvanted vaccine elicits >10^3^ geometric mean titer (GMT) of pseudovirus neutralizing antibodies that can be boosted with a second dose. These antibody levels are durable, with GMTs that surpass the post-prime levels for more than 5 months post-boost. Importantly, sera from these NHPs exhibit broad cross-reactivity against lentiviruses pseudotyped with spike proteins from MERS-CoV clades A, B, and C as well as a more distant pangolin merbecovirus. In transgenic mice expressing human DPP4, immunization provided dose-dependent protection against MERS-CoV lethal challenge, and in an established alpaca challenge model, immunization fully protected against MERS-CoV infection. This protein-based MERS-CoV nanoparticle vaccine is a promising candidate for advancement to clinical development to protect at-risk individuals and for future use in a potential outbreak setting.

## Introduction

The emergence of three human pathogenic betacoronaviruses since 2002 (SARS-CoV, MERS-CoV, and SARS-CoV-2) has demonstrated the substantial, continuing threat that coronaviruses pose to the human population. MERS-CoV, the causative agent of Middle East respiratory syndrome, was first identified in 2012 and has since caused 2613 laboratory-confirmed cases and 939 deaths (36% CFR)^1^. MERS-CoV likely originated in bats and is transmitted from its intermediate host, dromedary camels, to humans^2–5^. MERS-CoV undergoes human-to-human transmission, most notably in healthcare settings, and continues to be an ongoing threat as highlighted by five recent cases in 2024, four of which were fatal, in the Kingdom of Saudi Arabia^6,7^. There are currently no prophylactic vaccines or therapeutic interventions approved for use against MERS. The ongoing morbidity and mortality associated with MERS, the high prevalence of MERS-CoV in dromedary camel populations, and the potential for increased human-to-human transmission indicate a pressing need for a safe and effective MERS vaccine. The availability of such a vaccine would be useful as a routine prophylactic immunization for at-risk individuals such as healthcare workers, camel industry workers, and travelers to endemic regions. The vaccination of those at highest risk of MERS, due to occupation or travel, could mitigate the risk of a future pandemic and would establish the availability of a vaccine for broad use if it became needed.

Similar to SARS-CoV and SARS-CoV-2, MERS-CoV is a single-stranded RNA virus which displays a single surface protein, spike, responsible for host-cell recognition and viral entry^6,8^. MERS-CoV spike is a heavily glycosylated trimeric class I viral fusion protein that is expressed as a single polypeptide and cleaved via host cell proteases into S1 and S2 subunits, which remain non-covalently assembled following proteolysis^8^. The MERS-CoV spike receptor binding domain (RBD) in the S1 domain recognizes dipeptidyl peptidase 4 (DPP4)^9^ unlike SARS-CoV and SARS-CoV-2, both of which utilize angiotensin converting enzyme 2 (ACE2) as their receptor. Following binding, the virus is endocytosed, at which time a secondary proteolytic cleavage event occurs at the S2’ site within the S2 domain to allow viral membrane fusion to proceed^8^.

As with SARS-CoV-2, antibodies targeting the spike protein following MERS-CoV infection demonstrate neutralizing activity^10–12^. Thus, the spike protein has been an important target for experimental MERS vaccine candidates^13^. Several vaccine platforms, including viral-based, subunit, and DNA have been evaluated in preclinical studies, some of which elicited detectable MERS-CoV neutralizing antibodies and protected from viral challenge in preclinical models^14^. A few MERS-CoV vaccine candidates have been tested in early-stage clinical trials; however, no candidates have progressed to late-stage clinical development^15–19^. Therefore, advancing additional candidates towards clinical evaluation is critical.

Similar to other class I viral fusion proteins, the MERS-CoV spike trimer is a metastable protein poised to convert from the prefusion to the postfusion state upon receptor binding and cell entry^8^. Prior work has demonstrated that betacoronavirus spike proteins can be stabilized by mutating two key residues between the first heptad repeat (HR1) and second heptad repeat (HR2) to proline^20,21^. Installation of these two, as well as an additional four proline mutations in the SARS-CoV-2 spike increased protein expression and led to enhanced immunogenicity, presumably as a result of improved stability of the prefusion conformation^21,22^. We therefore based our MERS-CoV antigen design on a stabilized spike containing a mutated S1/S2 cleavage site and 3 proline mutations (Figure 1A). Structural studies of the MERS-CoV spike indicate that the region at the base of the trimer prior to the transmembrane domain (residues 1230-1294) is likely flexible, as it is unresolved via cryo-EM (Figure 1B)^23^. We thus hypothesized, based in part on our prior SARS-CoV-2 antigen design^24^, that removal of this flexible region may facilitate improved presentation of the spike in a conformation conducive to eliciting neutralizing antibodies. Furthermore, the eliminated region is a known epitope of non-neutralizing antibodies in the context of SARS-CoV- 2^25–27^, and we reasoned that it could be in MERS-CoV spike as well, further supporting the removal of this region.

**Figure 1.**
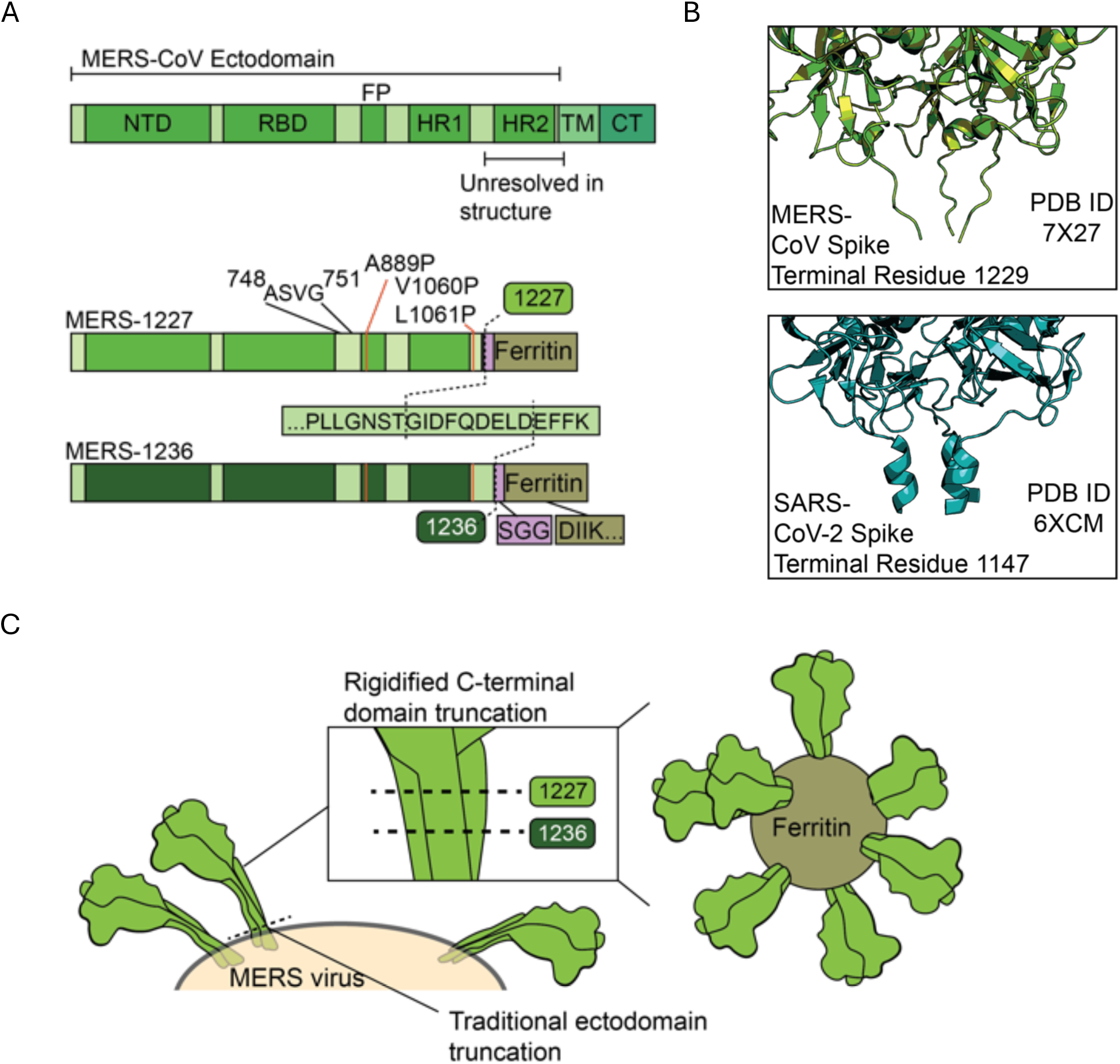
Structure-informed design of two novel MERS-CoV spike ferritin nanoparticle vaccine candidates. (A) Domain map of the MERS-CoV spike, containing the N-terminal domain (NTD), receptor-binding domain (RBD), fusion peptide (FP), heptad repeat 1 and 2 (HR1 and HR2), transmembrane domain (TM), and cytoplasmic tail (CT). MERS- 1227 and MERS-1236 contain a mutated furin cleavage site (ASVG) from residues 748-751, three stabilizing proline mutations (A889P, V1060P, L1061P), and ectodomain truncations after residue 1227 or 1236. MERS-1227 and MERS- 1236 encode a flexible linker (SGG) followed by *H. pylori* ferritin starting at residue 5. (B) Structural comparison of the MERS-CoV spike (top, PDB 7X27) with the SARS-CoV-2 spike (bottom, PDB 6XCM) reveals a flexible domain in MERS-CoV where a short helix is present in SARS-CoV-2. (C) Visual representation of the impact of the MERS-1227 and MERS-1236 ectodomain truncations on the presentation of spike on the surface of the ferritin nanoparticle.

Protein subunit vaccines have been shown in several contexts to have favorable stability profiles and to elicit robust, durable protection in humans^28–30^. In the absence of potent adjuvants, one limitation of protein subunit vaccines is weak immunogenicity compared to viral-based vaccines^29^. To overcome this limitation, proteins can be multimerized on nanoparticle platforms to confer benefits including facilitation of B cell cross-linking and enhanced neutralizing antibody responses^24,29,31–34^. Ferritin is a ubiquitous, naturally occurring, self-assembling protein that forms 24-subunit particles^35^. It contains a 3-fold axis of symmetry and is therefore an ideal platform for display of trimeric viral antigens like the MERS-CoV spike protein^31^. Ferritin nanoparticles have been demonstrated to be an effective and safe antigen multimerization tool in both preclinical and clinical studies^36–38^. We and others have shown that displaying the SARS-CoV-2 spike protein on the surface of *H. pylori* ferritin presents the antigen in a favorable conformation to elicit robust and cross-reactive neutralizing antibody responses^24,38–41^. Therefore, we applied these same antigen design principles to the MERS-CoV spike protein in an effort to develop a stable, safe, and effective vaccine for protection against MERS.

The results presented here comprise the design, biochemical characterization, stability profile, and immunogenicity assessment of novel MERS-CoV spike vaccine candidates. We demonstrate that our lead candidate, MERS-1227 ferritin nanoparticle (FNP), can be purified to homogeneity from a high-expressing Chinese hamster ovary (CHO) stable cell pool and retains antigenicity and nanoparticle structure following heat stress at 37 °C for 14 days. We show that MERS-1227 FNP is robustly immunogenic in BALB/c mice, NHPs, and alpacas as determined by MERS-CoV live-virus and pseudotyped lentiviral neutralization assays. In BALB/c mice expressing human DPP4, we observe dose-dependent protection from MERS-CoV lethal challenge following immunization with our lead MERS-CoV vaccine candidate. In an alpaca live-virus challenge model, we observed no detectable viral infection in alpacas immunized with Alhydrogel-adjuvanted MERS-CoV nanoparticles whereas 80% of the placebo-treated alpacas shed infectious virus following challenge. The robust stability, immunogenicity, and efficacy profile of this protein-based nanoparticle vaccine demonstrate optimal attributes for clinical development as a safe and efficacious vaccine against MERS.

## Results

### Design of two novel ferritin-based MERS-CoV spike vaccine candidates with varied ectodomain lengths

The MERS-CoV spike is a 1353 amino acid polypeptide that is cleaved into S1 and S2 subunits via proteolysis at residue 751 (Figure 1A)^8^. The ectodomain of MERS-CoV spike extends to residue 1294. We hypothesized that truncation of the ectodomain would facilitate more optimal presentation of the spike when displayed on the surface of a ferritin nanoparticle, potentially improving stability and immunogenicity. To determine the optimal truncation position for the spike ectodomain, we designed two constructs based on the spike sequence from MERS-CoV clade B England 1 strain^23^. These constructs extended through either residue 1227 (MERS-1227) or 1236 (MERS-1236) to optimally align the distance between the base of the protomers to the N-terminal residue of the ferritin 3-fold axis (Figure 1C). Following the ectodomain in both constructs, we included an SGG linker and residues 5-167 from *H. pylori* ferritin (Figure 1A) with an N19Q mutation in the ferritin sequence to eliminate a potential N-linked glycosylation site^31^. Additionally, we introduced three proline mutations guided by known SARS-CoV-2 2P and 6P variations that stabilize the spike protein^20–22^. We also mutated the S1/S2 furin cleavage site from RSVR to ASVG to prevent spike cleavage, as described in prior stabilized MERS-CoV constructs^20^.

### MERS-1227 and MERS-1236 can be purified to homogeneity from high-expressing CHO cell pools and form stable nanoparticles with expected antigenicity

To express and characterize large quantities of MERS-1227 and MERS-1236 FNPs, we generated stable, high-expressing cell pools at ATUM utilizing their proprietary Leap-In™ transposase technology and CHOK1-derived mi:CHO-GS™ host cells^42^. The cell line development process yielded two stable cell pools with protein titers of approximately 2.7 (MERS- 1227) and 3.2 (MERS-1236) grams per liter of cell culture in fed-batch supernatant harvested at day 14 as determined by SDS-PAGE quantitation (Figure S1A). We purified the nanoparticles at a research-level scale to a high degree of homogeneity using anion exchange chromatography in flow-through mode, dialysis, and size-exclusion chromatography as reported previously for a related ferritin nanoparticle vaccine candidate^40^. Given the simplicity of this process, we utilized materials purified in this manner for all subsequent analyses and immunizations described here. Additionally, we optimized a manufacturing process-applicable method for purification, which involved two bind and elute purification steps using an anion exchange membrane and a hydrophobic interaction chromatography (HIC) resin (Figure S1B). Both steps result in enrichment of the MERS-CoV FNP, demonstrating the feasibility of future at-scale process development of these complex macromolecules.

We characterized the purity of MERS-1227 and MERS-1236 FNPs from the research-scale purification using SDS-PAGE analysis (Figure 2A) and observed a pure species for both particles that migrates at the expected molecular weight of a MERS-CoV spike ferritin monomer (∼150 kDa). Both proteins exhibited a slight decrease in migration when analyzed under reducing conditions, likely due to disruption of intra-protomer disulfide bonds. We further analyzed the purity and the size of the purified FNPs using size-exclusion chromatography multi-angle light scattering (SEC-MALS). This analysis revealed highly homogenous particles, without notable high or low molecular weight species, and MALS-calculated molecular weights of 4.27 ± 0.05 MDa (MERS-1227) and 4.23 ± 0.09 MDa (MERS-1236) (Figure 2B). These observed molecular weights are slightly larger than those calculated from the MERS-1227 and MERS-1236 primary amino acid sequences (3.6 and 3.7 MDa, respectively). We hypothesized that this difference was due to glycosylation of the MERS-CoV spike, which has previously been characterized^43–45^. To evaluate glycosylation of our MERS-CoV FNP proteins, we performed mass spectrometry-based peptide mapping. We observed extensive N-glycosylation and limited but detectable O-glycosylation throughout the MERS-1227 and MERS-1236 spikes but not within the ferritin nanoparticle core (Figure S2 and Table S1), consistent with prior analyses of purified MERS-CoV spike proteins^43–45^.

**Figure 2.**
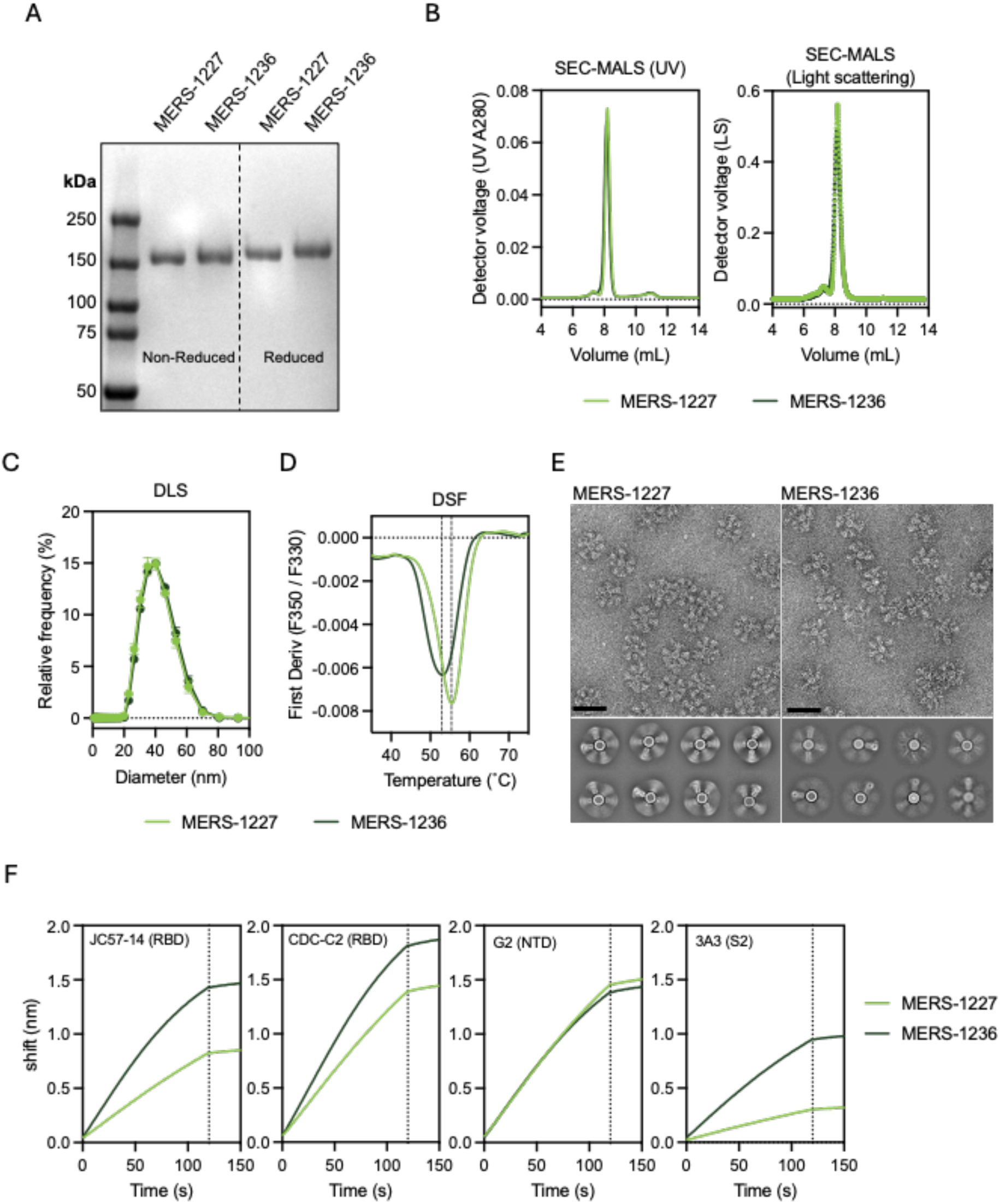
MERS 1227 and MERS-1236 nanopartiicles are comparablie in size and homogeneity but have distinct stability, structural, and antigenicity features. (A) SOS-PAGE analysis of MERS-1227 and IMIERS-1I236 purified from 14-day fed-batch supematants from stable CHO oell pools. (B) SEC-IMAlS reveals that purified MERS-1227 and MERS-1236 are homogenous nanopartic:les, with minimal aggregate species observed to the left of the nanopartic:le peak and no lower molecular weight species observed to the ri.ght (C} DLS analysis of IMIERS-1227 and MERS-1236 reveals a particle di,ameter of -40 nm. (D) IDSF shows that MERS-1236 has a melting1temperature -4 ·c lower than MIERS-1227 and a broader melting transition. (E) TEM micmgraphs (left) and 2D cl.ass averages (right) o-f MERS-1227 and IMIERS-1236 indicate that the spl1ke on the surface of MERS-1236 is more flexible compared to MERS-1227. Scale bar below micrographs represen,ts 50 nm. (F) BLII binding of MERS CoV spike-specific mAbs to IMIERS-1227 and MERS-1236.

Additional size characterization via dynamic light scattering (DLS) showed a diameter of 38.1 nm for MERS-1227 and 39.1 nm for MERS-1236 (Figure 2C), consistent with the expected 10 nm diameter of an *H. pylori* ferritin nanoparticle^35^ with two ∼14 nm MERS-CoV spikes projecting radially^8^. Interestingly, differential scanning fluorimetry (DSF) revealed distinct melting temperatures of 55.4 °C for MERS-1227 and 52.9 °C for MERS-1236, with MERS-1236 also exhibiting a broader melting profile (Figure 2D). This result suggests that the 9 additional amino acids at the C-terminus of the ectodomain of MERS-1236 influence protein stability. However, the most striking difference between MERS-1227 and MERS-1236 was observed by negative stain transmission electron microscopy (TEM) (Figure 2E), which showed that MERS-1236 spikes exhibit dramatically more structural flexibility. Collectively, these biochemical and structural data demonstrate that MERS-1227 displays a more rigid, stable functionalized nanoparticle, projecting the MERS-CoV spike protein radially from the ferritin core. Based on comparison of MERS-1227 and MERS-1236 via TEM and DSF, we elected to prioritize preclinical characterization of MERS-1227 and included MERS-1236 as a comparator when necessary. The structural and stability advantages of MERS-1227 suggested to us that it may exhibit fewer liabilities as a candidate for vaccine development.

To evaluate the antigenicity of MERS-1227 and MERS-1236, we utilized a panel of previously characterized monoclonal antibodies (mAbs) that bind to distinct epitopes across the MERS-CoV spike and used biolayer interferometry (BLI) to measure their binding to the nanoparticles. Within this panel, we included two RBD antibodies (JC57-14 and CDC-C2)^12^, an NTD antibody (G2)^46,47^, and an S2 antibody that binds an internal trimer epitope (3A3)^48,49^ (sequences provided in Table S2). As shown in Figure 2F, subtle differences in the antibody binding profiles of MERS-1227 and MERS-1236 suggest that epitopes on the spike are presented differently on these particles. Specifically, MERS-1236 showed moderately enhanced binding to the two RBD antibodies, which are both known to bind the MERS-CoV RBD in the “up” conformation^8,12^. This could suggest that this RBD conformation is favored in MERS-1236 as compared to MERS-1227. The NTD antibody G2 shows equivalent binding to both nanoparticles, and the internal S2-directed antibody 3A3 binds more strongly to MERS-1236. The 3A3 antibody binds at the trimer interface which is only accessible in an open conformation of SARS-CoV-2 spike that the trimer reversibly samples^48^. The increased binding of MERS-1236 to 3A3 is therefore consistent with a more flexible trimer, also supported by the TEM comparison of MERS- 1227 and MERS-1236 (Figure 2E) and may suggest MERS-1236 samples an “open trimer” conformation more frequently.

### MERS-1227 and MERS-1236 are stable over a range of pH and temperature conditions

Low-pH treatment is commonly employed for viral inactivation during downstream process development, and knowledge of pH tolerance is helpful in optimizing chromatography steps. Therefore, we sought to understand the effect of low pH on MERS-1227 and MERS-1236 nanoparticle stability and antigenicity. We formulated purified MERS-1227 and MERS-1236 in pH 7.5 formulation buffer and held them in the pH-adjusted solutions at 2, 3, 3.5, and 4 for 2 hours.

Then, the nanoparticles were dialyzed back into pH 7.5 buffer and evaluated using analytical SEC, DLS, DSF, and BLI (Figure 3A-B). These analyses revealed that pH 2 treatment causes irreversible damage to both MERS-1227 and MERS-1236 as observed by all biophysical measurements, suggesting dissociation of the nanoparticle and/or denaturation of the MERS-CoV spike. BLI revealed that binding of an RBD mAb (CDC-C2) and an NTD mAb (G2) is retained following pH 2 treatment, which may suggest that antigenicity of the isolated protomer is retained to some degree following nanoparticle dissociation. Treatments at pH 3, 3.5, and 4 do not notably affect particle structure or antigenicity.

**Figure 3.**
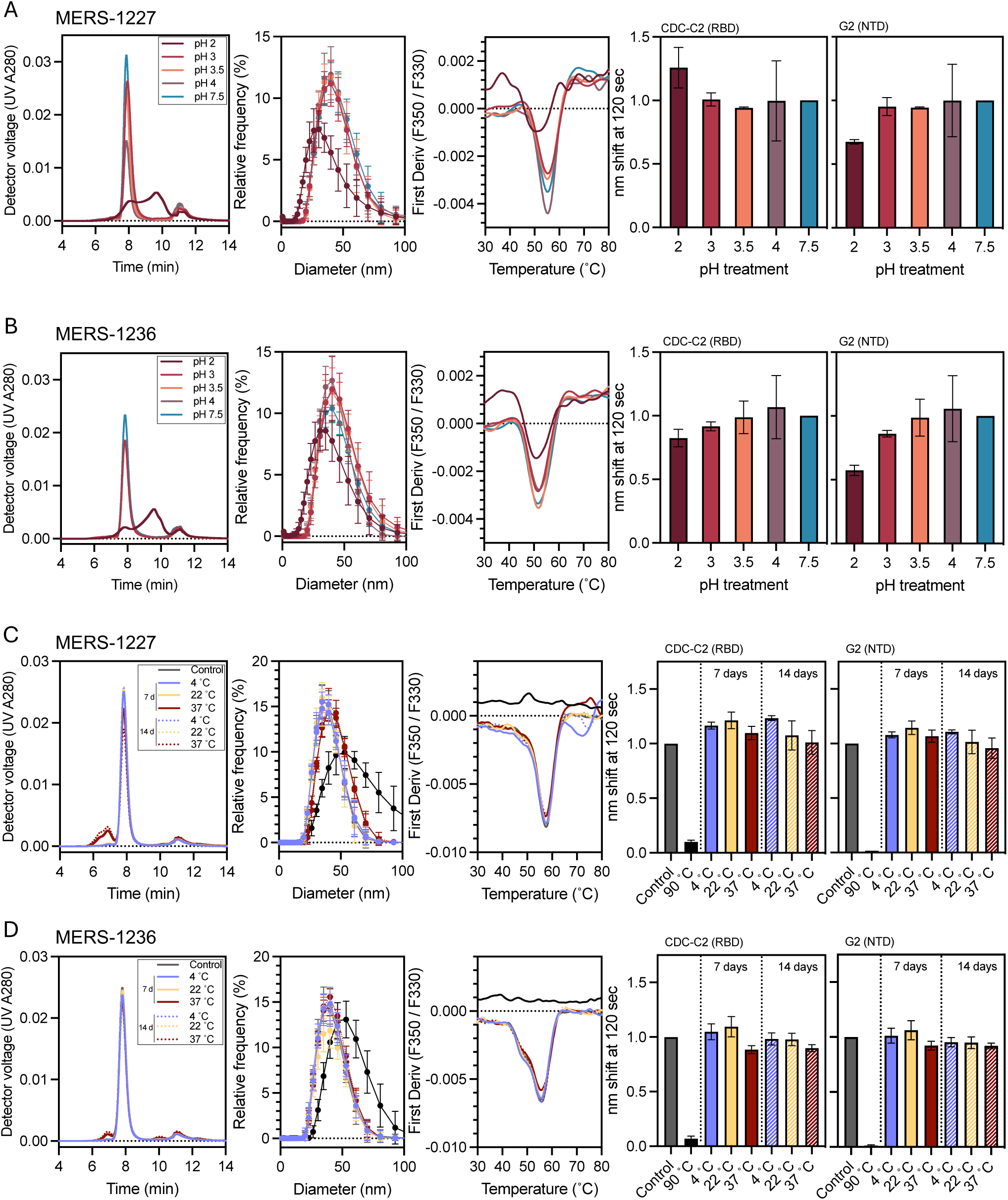
MERS-1227 and MERS-1236 retain nanoparticle structure and antigenicity following low-pH treatment and extended incubation up to 37 °C. (A) Biophysical characterization of MERS-1227 following pH treatment at pH 2, 3, 3.5, 4, and 7.5. Panels correspond to analytical SEC (left), DLS (middle left), DSF (middle right), and BLI (right two panels) with an RBD-targeting mAb, CDC-C2, and an NTD-targeting antibody, G2. pH treatments were conducted in duplicate and each sample was then characterized using each assay. Curves shown for SEC, DLS, and DSF represent one replicate. Error bars on the DLS plot represent standard deviation. BLI plots show the mean nm shift value of the two treatment replicates at 120 s normalized to the pH 7.5 control, and error bars represent SD. (B) Biophysical characterization of MERS-1236 following pH treatment as described in (A). (C) Biophysical characterization of MERS- 1227 following incubation at 4, 22, or 37 °C for 7 and 14 days. Solid lines represent 7-day treated samples and dashed lines represent 14-day treated samples. Control samples include a sample not subjected to any incubation (gray) and a sample heat-treated at 90 °C for 30 min (black). The 90 °C heat-treated sample was included in the DLS, DSF, and BLI analyses but was excluded from SEC analysis. BLI measurements were normalized to the untreated control sample. (D) Biophysical characterization of MERS-1236 following thermal incubations as described in (C).

Temperature stability is important in vaccine development with the goal of optimizing a modality and formulation such that cold-chain requirements can be minimized. This is especially relevant to a MERS vaccine, as the endemic regions where it would be distributed regularly experience high ambient temperatures. To understand the thermostability of MERS-1227 and MERS-1236, we formulated purified materials in formulation buffer and stored them at 4, 22, or 37 °C for 14 days. Following these temperature holds, we evaluated the quality and stability of the FNPs using analytical SEC, DLS, DSF, and BLI (Figure 3C-D). We confirmed that DLS, DSF, and BLI binding to these mAbs are appropriate metrics of protein integrity, as incubation for 30 minutes at 90 °C notably perturbed particle size (DLS) and fully ablated the melting curve (DSF) and binding to both mAbs (BLI) for both FNPs (Figure 3C-D). By SEC analysis, we observed that both MERS-1227 and MERS-1236 remained stably associated as nanoparticles even after 14 days at 37 °C, albeit with a minor fraction of particles forming an aggregate species as evidenced by a peak at ∼7.0 min that elutes prior to the nanoparticle peak at ∼8.0 min. DLS showed no major change in the size of particles following heat-treatment, and DSF revealed no change in the primary melting peak following the temperature hold conditions, providing evidence for the conformational stability of the particles. We also found that both MERS-1227 and MERS-1236 retain antigenicity against an RBD-specific (CDC-C2) and an NTD-specific (G2) monoclonal antibody after 14 days at either 22 or 37 °C (Figure 3C-D).

*MERS-1227 and MERS-1236 are more immunogenic compared to trimer alone in BALB/c mice following one or two doses*.

We sought to determine the effect of spike multimerization on elicitation of neutralizing antibodies by immunizing BALB/c mice (n = 5 or 10) with 2 or 10 µg of MERS-CoV full-length spike ectodomain (residues 1-1294) with a GCN4 trimerization domain^50^ or 0.4, 2, or 10 µg of MERS-1227 or MERS-1236 adjuvanted with 150 µg Alhydrogel (Figure 4A). We collected serum at day 21 (following a single dose) and day 42 (following two doses). As seen in Figure 4B, after a single dose, mice immunized with any dose level (0.4, 2, or 10 µg) of either MERS-1227 or MERS-1236 FNPs elicited ∼2- to 3-fold higher levels of both anti-spike and RBD IgG antibodies as compared to mice immunized with MERS-CoV trimer. Following a second dose, mice across the three antigen groups at all doses have similar levels of MERS-CoV spike and RBD IgG (Figure 4B, right panels).

**Figure 4.**
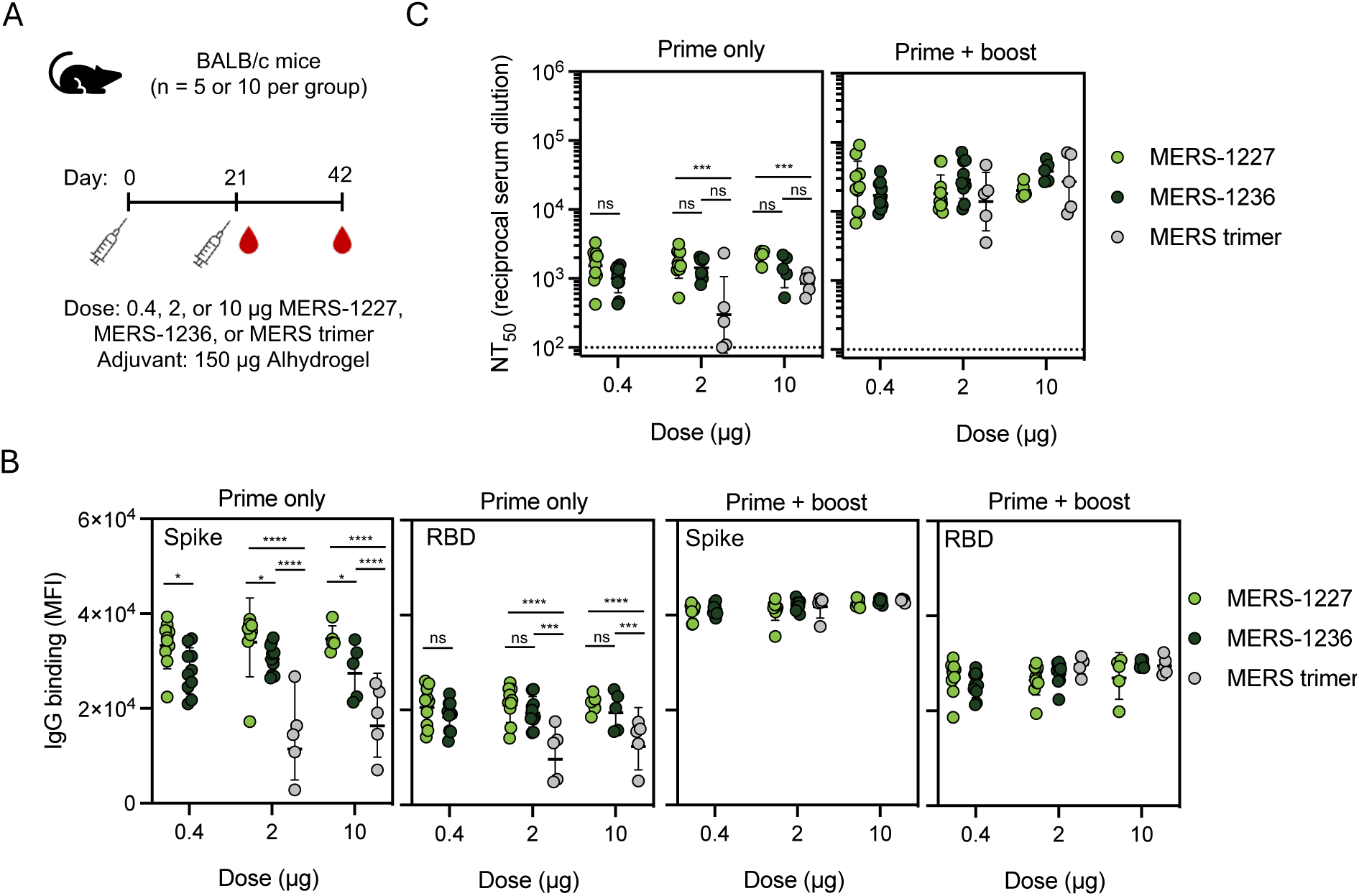
Immunization of BALB/c mice with MERS-1227 or MERS-1236 results in more robust binding and neutralizing antibody responses compared to soluble trimer. (A) Immunization schedule. For MERS- 1227 and MERS-1236, mice were immunized with either 0.4 (n = 10), 2 (n = 10), or 10 µg (n = 5) FNP. For MERS-CoV soluble trimer, mice were immunized with either 2 (n = 5) or 10 µg (n = 5). All doses were formulated with 150 µg Alhydrogel. Serum was collected at study day 21 and 42. (B) Antigen-specific IgG responses from each mouse measured at 1:100 dilution using a Luminex assay. Luminex beads coated with soluble MERS-CoV trimer or MERS-CoV RBD were used to evaluate levels of binding antibodies. Each dot represents the average of the mean fluorescence intensity (MFI) value per mouse, lines represent geometric mean, and error bars represent the geometric SD. (C) MERS-CoV spike-pseudotyped lentivirus neutralization GMTs observed following a prime only (day 21, left panel) and a prime + boost (day 42, right panel). Each circle represents the NT_50_ value for an individual mouse obtained by running pseudovirus neutralization analysis in quadruplicate (with the exception of one mouse in the 10 µg trimer group at day 21 in duplicate). Lines represent the GMT, and error bars represent the geometric SD. Statistical analysis in (B) and (C) was conducted using a two-way ANOVA with multiple comparisons. No statistical differences within dose groups were observed at the prime + boost timepoint. ****, p < 0.0001; ***, p < 0.001; **, p < 0.01; *, p < 0.05.

We quantified neutralizing antibody titers using a luciferase-based assay^51^ with lentivirus pseudotyped with MERS-CoV spike and HeLa cells stably expressing human DPP4 as the targets of infection. We validated our assay using MERS-CoV specific neutralizing antibodies with known IC50 values^12,52^. The values obtained in our assay concorded well with those reported in the literature (Figure S3A). As expected, no neutralization was detected with a negative control SARS-CoV-2 specific mAb, casirivimab. We also used convalescent serum from a camel challenged with MERS-CoV with a known PRNT90 of 1:320 serum dilution as a benchmark for our pseudovirus neutralizing titers (Figure S3B)^53^. In our pseudovirus assay, this camel serum sample inhibits 90% of viral entry (IC90) at a serum dilution of 1:6810. This result indicates that our lentivirus-based pseudovirus assay exhibits higher sensitivity than a plaque-reduction neutralization assay using live MERS-CoV virus, as reported for other lentivirus-based neutralization assays compared to live viral comparators^54^.

Following a single dose, we observed robust pseudovirus neutralizing titers of GMT ∼10^3^ for MERS-1227 and MERS-1236 at all doses tested, suggesting that even 0.4 µg (when adjuvanted with 150 µg Alhydrogel) is a saturating dose in BALB/c mice (Figure 4C, left panel). Comparatively, MERS trimer elicited a lower and more heterogeneous antibody response at 2 µg and a more homogenous but still overall lower response at 10 µg. After a second dose, we observed a boost in titers across all antigens and dose groups. Notably, the MERS-CoV trimer groups show a higher degree of variability, especially at the 10-µg dose level, with a geometric standard deviation of 2.6 as compared to 1.3 and 1.4 for MERS-1227 and MERS-1236, respectively (Figure 4C, right panel). We additionally tested pools of serum from the MERS-1227 and MERS-1236 groups and confirmed they neutralized live MERS-CoV in a plaque-reduction neutralization assay (Table S3). This initial mouse immunogenicity study revealed that the multimerized, stabilized MERS-CoV spike elicited a stronger neutralizing antibody response following one dose and a more homogeneous response following two doses compared to soluble trimer. Subsequently, we performed an additional dose de-escalation study with Alhydrogel-adjuvanted MERS-1227 and found that it elicited a neutralizing antibody response at 0.016 µg, the lowest dose tested, following a prime and a boost (Figure S4).

### MERS-1227 and MERS-1236 elicit robust neutralizing antibody responses in NHPs following a single dose that increase and persist at least several months following a second dose

To evaluate MERS-1227 and MERS-1236 immunogenicity in an NHP model, we immunized cynomolgus macaques (n = 5 per antigen) with 50 µg FNP adjuvanted with 750 µg Alhydrogel and measured the magnitude, durability, and breadth of the antibody response following one and two doses (Figure 5). Following a single immunization, we observed pseudovirus neutralizing antibody titers at or greater than reciprocal serum dilution (NT50) of 10^3^ for all animals in both vaccine groups (Figure 5B). Importantly, we did not observe notable waning of these titers over the three-month period following the prime vaccination. We immunized the NHPs with a homologous boost at study day 93 and observed a robust increase of greater than 10-fold in pseudovirus neutralizing titers at study day 100 that continued to increase to study day 127. At study day 253 (160 days post-boost), titers remained higher than those observed following a single dose. Importantly, the rate of decay observed between days 180 and 253 was decreased compared to that observed between days 127 and 180, which is suggestive of the potential for a durable response as has been reported for other alum-adjuvanted protein-based vaccines^55^. To further characterize the serum response, we used a Luminex panel containing MERS-CoV spike, RBD, and NTD, and evaluated total IgG levels to these antigens. We observed similar spike and RBD IgG levels, with slightly more waning of the spike titers observed between study days 188 and 253. Interestingly, spike and RBD binding titers appear to decrease less quickly than pseudovirus neutralizing titers (Figure 5, compare B to C). Overall, NTD antibody levels were lower than spike or RBD levels, suggesting that the RBD is a more immunodominant target of these antigens.

**Figure 5.**
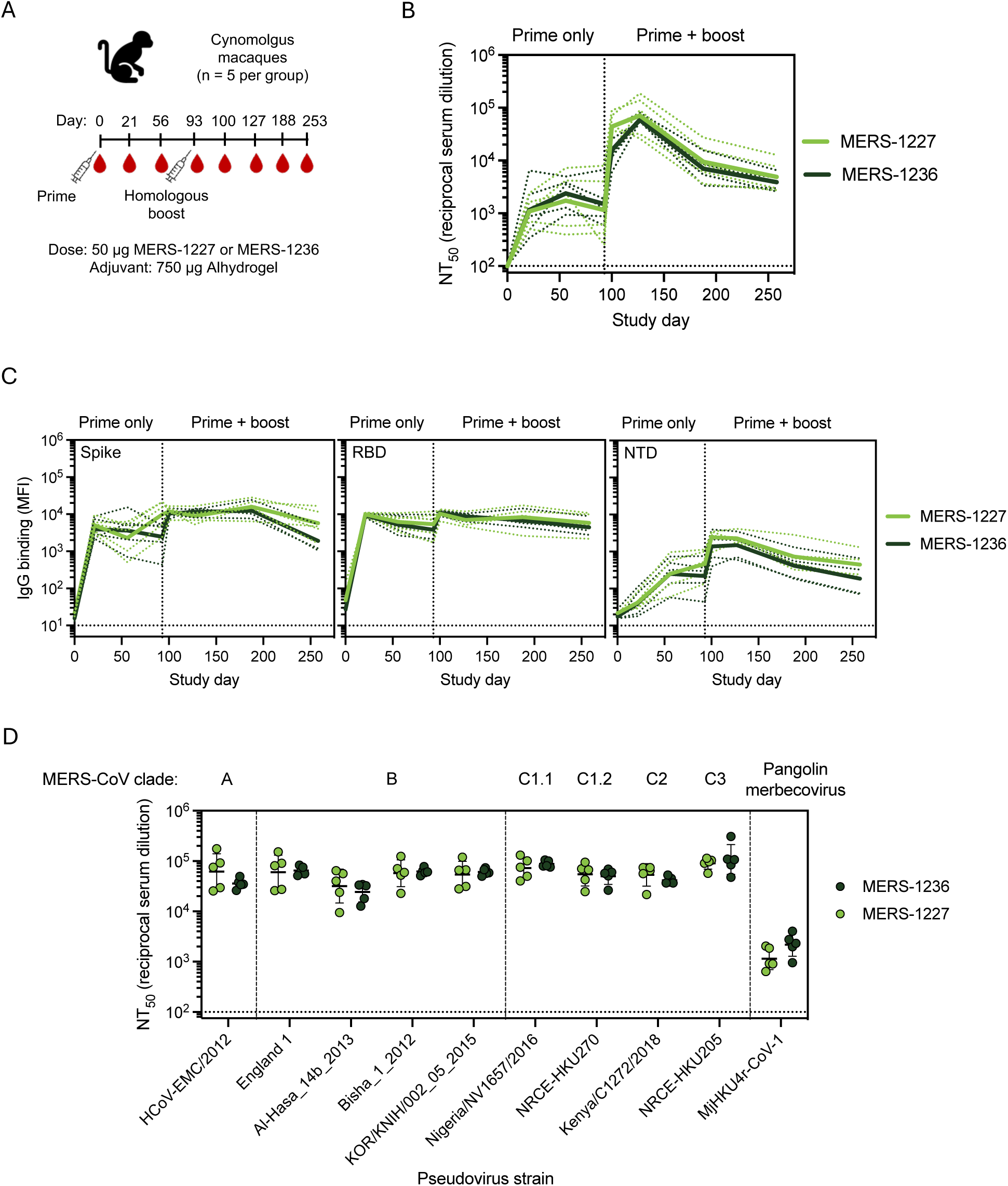
NHPs immunized with Alhydrogel-adjuvanted MERS-1227 and MERS-1236 elicit robust, broad, and durable neutralizing antibody responses. (A) NHP immunization schedule with serum collection on days indicated by a blood drop symbol. (B) MERS-CoV spike pseudotyped lentivirus neutralization GMTs following a prime and a boost. The solid lines represent the average GMTs of each antigen group and the dashed lines represent the NT_50_ values by individual NHP. (C) Antigen-specific IgG responses from immunized NHPs measured at 1:100 dilution using a Luminex assay. Luminex beads coated with either soluble MERS-CoV trimer (left), MERS-CoV RBD (middle), or MERS-CoV NTD (right) were used to evaluate levels of binding antibodies. The solid line represents the average GMTs per antigen group, and the dashed lines represent the MFI values per individual NHP. (D) NHP serum collected ∼5 weeks following a second immunization (study day 127) neutralizes a panel of lentiviruses pseudotyped with spike sequences from MERS-CoV clades A, B, C, and a pangolin merbecovirus (MjHKU4r-CoV-1). Each circle represents the NT_50_ value for an individual NHP obtained by running pseudovirus neutralization analysis in quadruplicate. Lines represent the GMT and error bars represent the geometric SD.

To characterize the breadth of response to known MERS-CoV strains, we established a panel of MERS-CoV spike-pseudotyped lentiviruses comprising clades A, B (currently circulating in humans and the vaccine strain in this study), C and a distantly related pangolin merbecovirus, MjHKU4r-CoV-1^56,57^. We observed robust neutralization of all MERS-CoV clades in the panel (Figure 5D), including clade C strains which are known to circulate in camels but not humans^56^. Remarkably, we also observed neutralization of MjHKU4r-CoV-1 which only shares a 66% amino acid sequence identity with the spike of England 1 MERS-CoV, albeit at reduced potency relative to other strains tested^57^. We did not observe any neutralization of the MjHKU4r-CoV-1 pseudotyped lentivirus with pre-immune NHP serum (Figure S5). Pseudovirus neutralization, breadth, and total IgG levels were highly similar between MERS-1227 and MERS-1236 immunized NHPs. Overall, the results of this immunization study demonstrate that adjuvanted MERS-1227 and MERS-1236 elicit a robust, durable, and broad response in NHPs that can be boosted following a second dose administered at a 3-month interval, which is consistent with other effective vaccine regimens^58^.

### The immune response to MERS-1227 can be tuned using different adjuvant combinations

While alum is the most widely used vaccine adjuvant with a long clinical history, alternative adjuvants have potential to enhance the immunogenicity and durability of protein subunit vaccines^59^. We therefore formulated MERS-1227 with a panel of adjuvants known to engage distinct immune pathways^59^, including CpG (ODN2395), Quil-A + monophosphoryl lipid A (MPLA), AddaVax, and 3M-052 and probed the resulting immunogenicity in mice. Based on prior preclinical and clinical use of these adjuvants, we evaluated some co-formulated with Alhydrogel (Quil-A+MPLA, CpG, and 3M-052) as well as some without Alhydrogel (AddaVax, Quil-A+MPLA, and 3M-052). We formulated MERS-1227 with the various adjuvants and immunized BALB/c mice with a 0.4 µg dose of FNP protein followed by a homologous boost at day 21 (Figure 6A). We collected serum at day 21 (prime only) and three weeks following the boost dose at day 42 (prime + boost) to assess the elicitation of neutralizing antibodies.

**Figure 6.**
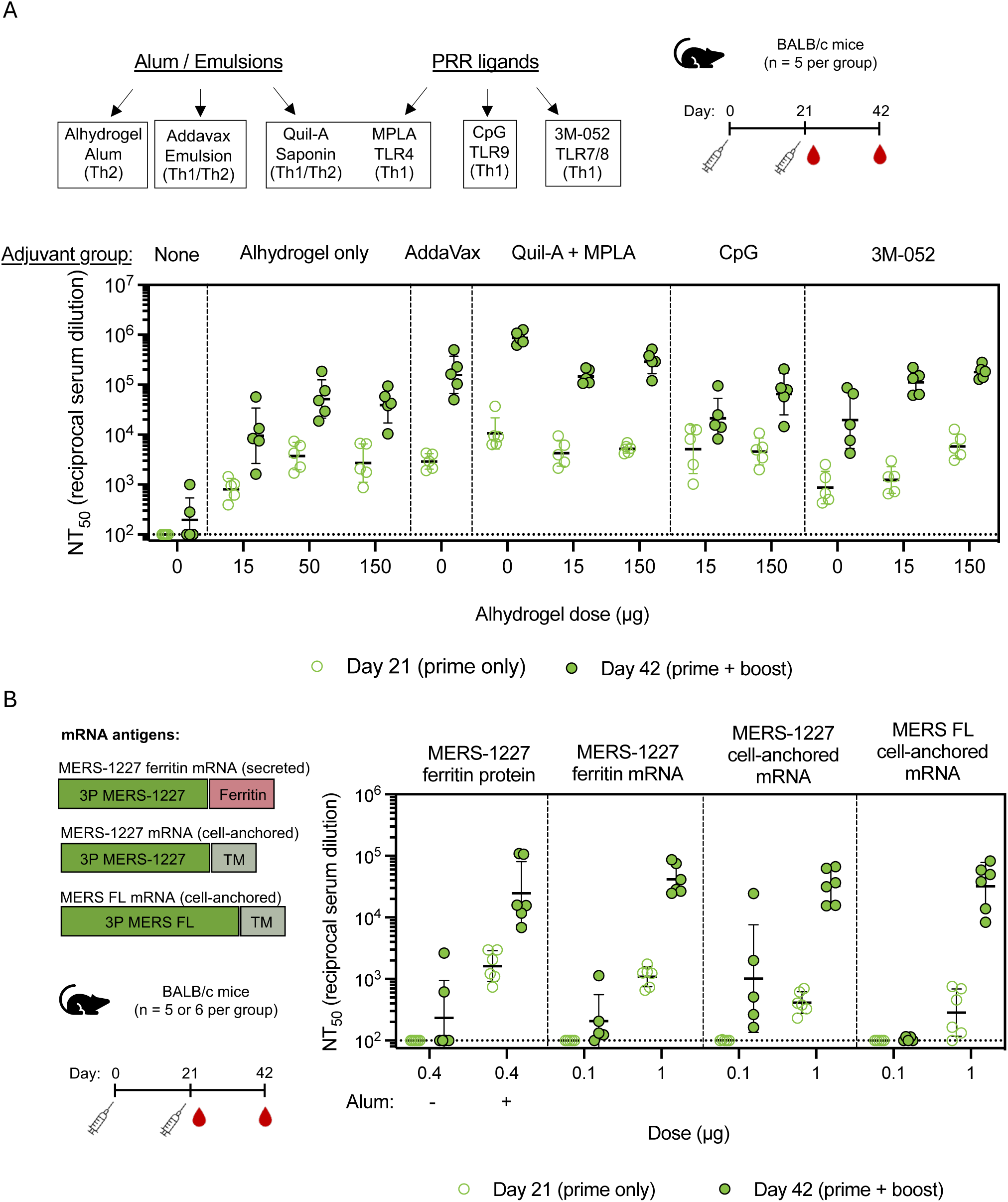
Evaluation of MERS-1227 FNP protein immunized with varied adjuvants and MERS-1227 encoded as an FNP by mRNA reveals optionality for formulation and modality. (A) Adjuvants evaluated included a range of alum/emulsion and PRR agonists. Mice (n = 5 per adjuvant group) were immunized at day 0 and 21 with 0.4 µg MERS- 1227 FNP protein, and serum was collected at day 21 (open circles) and 42 (closed circles). Each circle represents the NT_50_ value for an individual mouse obtained by performing pseudovirus neutralization analysis in quadruplicate. Lines represent the GMT, and error bars represent the geometric SD. (B) mRNA antigens evaluated for immunogenicity included mRNA-encoded MERS-1227 ferritin, cell-anchored MERS-CoV spike containing a deletion of ectodomain residues 1228-1294, and cell-anchored MERS-CoV spike with a full-length ectodomain. Mice (n = 5 or 6 per group) were immunized at day 0 and 21 with either 0.4 µg MERS-1227 FNP (n = 6), 0.1 µg mRNA (n = 5), or 1 µg mRNA (n = 6). Serum was collected at day 21 (open circles) and 42 (closed circles). Each circle represents the NT_50_ value for an individual mouse obtained in quadruplicate. Lines represent the GMT, and error bars represent the geometric SD.

An adjuvant is required to elicit a robust response as unadjuvanted MERS-1227 generated markedly lower pseudovirus neutralization titers compared to all three doses of Alhydrogel tested (Figure 6A). Titration of Alhydrogel in the absence of other adjuvants indicated that in mice, the dose response is maximal between 15 and 50 µg, with no improvement when comparing 50 to 150 µg. Immunization with MERS-1227 formulated with AddaVax (similar to MF59^60^ in clinically approved vaccines) did not improve titers compared to the mid- and high-dose Alhydrogel groups following a single dose but did show a stark improvement after the boost.

The most striking improvement in immunogenicity was observed with Quil-A+MPLA, especially after two doses. While addition of Quil-A+MPLA to Alhydrogel improved responses compared to the matched 15 µg and 150 µg Alhydrogel-only groups, the strongest response to Quil-A+MPLA was observed in the absence of Alhydrogel. Addition of CpG (ODN2395) only conferred an immunogenicity improvement compared to Alhydrogel alone at the 15 µg Alhydrogel dose, with minimal difference between the 150 µg Alhydrogel +/- CpG groups at both timepoints. Interestingly, addition of 3M-052, a lipidated TLR7/8 agonist^61^, did not result in increased neutralizing titers compared to the Alhydrogel only dose groups after a single dose but did show an improved response following a boost, most notably in the presence of either 15 or 150 µg Alhydrogel.

To evaluate the influence of adjuvants on the ratio of Th1/Th2 responses following immunization with MERS-1227, we bound biotinylated MERS-CoV spike trimer to streptavidin Luminex beads and quantified the levels of mouse IgG2a and IgG1 (Figure S6). In mice, induction of IgG1 is associated with a Th2-biased response and IgG2a is associated with a Th1-biased response^62^. Mice immunized with MERS-1227 adjuvanted with either 15, 50, or 150 µg Alhydrogel alone elicited an IgG response strongly dominated by IgG1, indicative of a Th2 response, which is well-documented for alum adjuvants^63^. AddaVax was observed to have a slightly decreased Th2-skewing as compared to Alhydrogel alone. Importantly, we observed that co-formulation of Alhydrogel with either Quil-A+MPLA or 3M-052 generated a Th1 bias characterized by a higher IgG2a / IgG1 ratio in these groups compared to Alhydrogel alone. CpG was observed to also overcome the Alhydrogel-induced Th2 skew at the 15 µg Alhydrogel dose, but not at the 150 µg dose. Taken together, the results of this adjuvant screen demonstrate several feasible mechanisms to improve both humoral and cell-mediated immune responses to MERS-1227.

### MERS-1227 elicits a comparable response when administered as an adjuvanted protein nanoparticle or an mRNA-encoded ferritin nanoparticle

The remarkable success of the SARS-CoV-2 mRNA vaccines has demonstrated the feasibility of widespread immunization using mRNA platforms. While protein vaccines exhibit advantages relating to thermostability and lower cost-of-goods, mRNA technology is advancing to meet these challenges and can be produced and modified quickly if needed to respond to a novel or mutated pathogen^64^. For this reason, we evaluated immunization using mRNAs packaged in SM102-containing lipid nanoparticles (LNPs)^65^ encoding either MERS-1227 ferritin, a cell-anchored form of the MERS spike containing a deletion of the ectodomain from residue 1228-1294, or a full-length MERS spike with no ectodomain deletion (Figure 6B). All constructs contained 3P stabilizing mutations, a mutated furin cleavage site, and a 21-residue deletion at the C-terminus of the protein to remove the cytoplasmic tail. We immunized naïve BALB/c mice with either 0.1 µg or 1 µg mRNA on day 0 and day 21 and collected serum at days 21 (prime only) and 42 (prime + boost) (Figure 6B). As a comparator, we included the MERS-1227 FNP protein +/- 150 µg Alhydrogel.

Following a single dose, both the Alhydrogel-adjuvanted MERS-1227 FNP protein and MERS-1227 FNP encoded on mRNA (1 µg dose) elicited similar responses that were higher than both cell-anchored forms of the MERS spike. After boosting, the 1 µg mRNA groups all exhibited similar levels of pseudovirus neutralizing titers, which were comparable to the adjuvanted MERS- 1227 protein nanoparticle. Interestingly, the 0.1 µg mRNA dose groups elicited minimal responses after both doses, and the only responders at this dose level were seen in the MERS-1227 ferritin and the MERS-1227 cell-anchored mRNA groups. This suggests that the truncation of the ectodomain to remove the flexible region between 1228 and 1294 conferred a benefit in neutralizing response in the mRNA format, which could provide important insight into the design of other mRNA antigens for vaccine delivery. Furthermore, these results show the versatility of the 3P stabilized spike we have designed to be administered as a protein nanoparticle, an mRNA-encoded protein nanoparticle, and as a cell-anchored stabilized spike.

### Immunization with MERS-1227 provides dose-dependent protection in hDPP4 BALB/c mice

To evaluate efficacy of MERS-1227 in a small animal model, we utilized a BALB/c mouse model in which the human DPP4 receptor is expressed in the nasal turbinates, trachea, lungs, and kidney^66^. We immunized hDPP4 transgenic mice (n = 8 per group) with 0, 0.016, or 0.4 µg MERS-1227 formulated with 150 µg Alhydrogel at days 0 and 21 and collected serum at days 21 and 42 (Figure 7A). MERS-CoV pseudovirus and live-virus neutralization titers (Figure 7B and 7C) showed dose-dependence, and pseudovirus titers were consistent with our evaluation in wild-type BALB/c mice (Figure S4).

**Figure 7.**
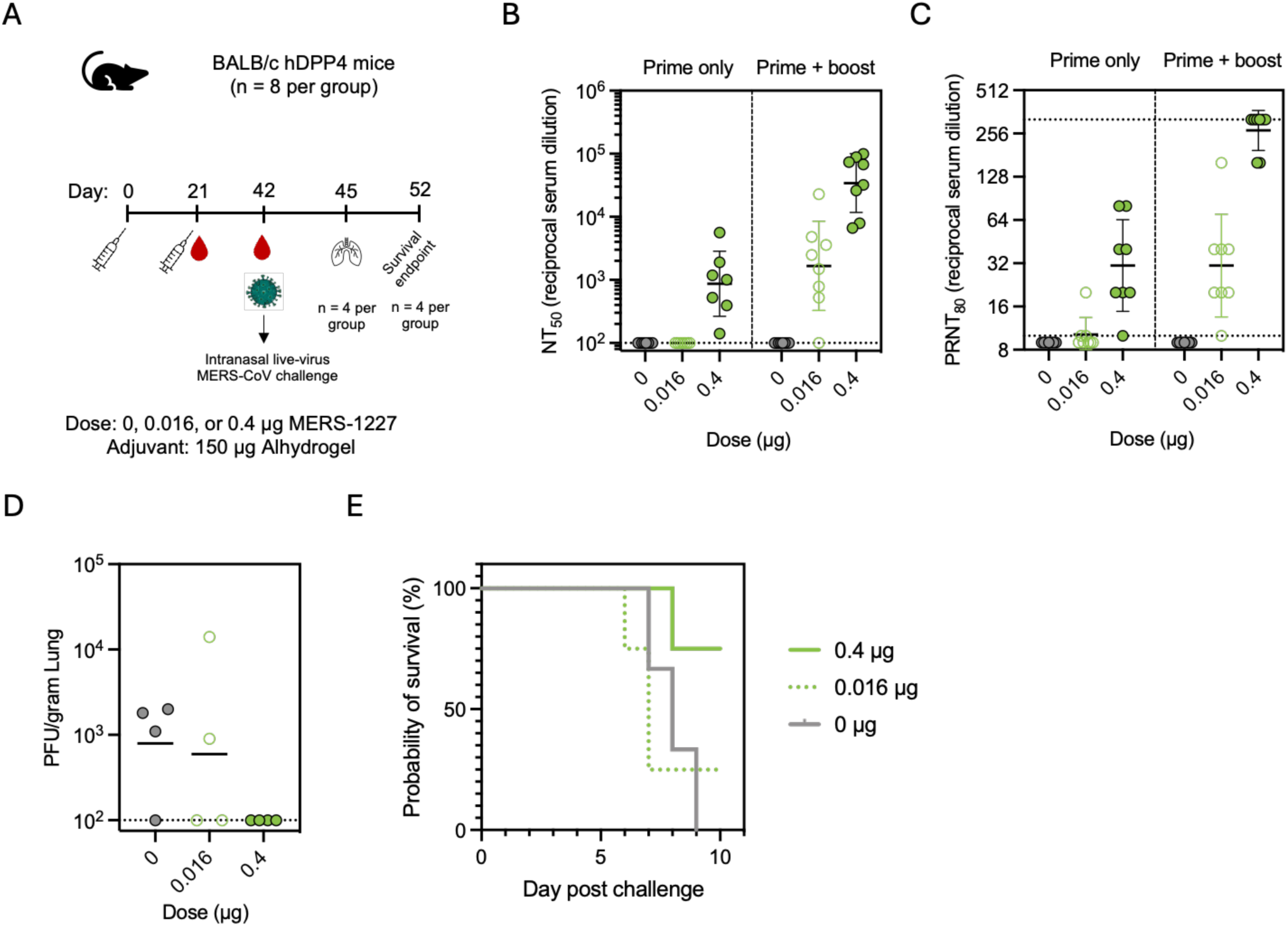
Immunization with MERS-1227 protects human DPP4 transgenic mice from lethal MERS-CoV **challenge.** (A) hDPP4 mouse challenge study design and timeline. Mice were challenged via intranasal inoculation with EMC/2012 MERS-CoV. (B) Pseudovirus neutralizing titers at study day 21 (prime only) and 42 (prime + boost). Each circle represents the NT_50_ value for an individual mouse obtained by running pseudovirus neutralization analysis in quadruplicate. Serum from one mouse in the 0.016 µg and one mouse in the 0.4 µg dose groups were not analyzed for day 21 neutralization due to lack of available serum. Lines represent the GMT and error bars represent the geometric SD. (C) Live-virus neutralization obtained from a plaque reduction neutralization assay using day 21 (prime only) and 42 (prime + boost) serum. Each circle represents the PRNT_8Ũ_ value for an individual mouse. Lines represent GMT and error bars represent geometric SD. (D) Quantitation of live MERS-CoV virus in lung samples obtained from mice sacrificed 3 days post-challenge. (E) Survival curves post challenge. One mouse in the 0 µg group died pre-challenge and is not shown in the survival curve. The mouse in the 0.4 µg dose group that succumbed to disease on day 8 was determined to be pregnant at necropsy.

Three weeks following the boost dose, mice were challenged via intranasal inoculation with 10,000 PFU of the EMC/2012 strain of MERS-CoV. Four mice from each group were sacrificed 3 days post-challenge and live-viral titers of MERS-CoV were quantified from the lungs of these animals using plaque assays (Figure 7D). Three mice in the placebo group had viral titers (ý 0.9 x 10^3^ PFU / gram lung) in the lung as compared to two mice in the 0.016 µg group and no mice in the 0.4 µg group, demonstrating protection from viremia following MERS-1227 vaccination. The remaining four mice in each group were monitored for weight loss (Figure S7) and survival (Figure 7F) until day 10 post-challenge. All mice in the placebo group succumbed to challenge by day 9, whereas 1/4 mice in the 0.016 µg group and 3/4 mice in the 0.4 µg group survived to day 10. Of note, the mouse that died in the 0.4 µg group prior to day 10 was found to be pregnant during necropsy. Taken together, the lung viral titer and survival results post viral challenge indicate that immunization with Alhydrogel-adjuvanted MERS-1227 was protective against lethal MERS-CoV challenge in hDPP4 mice in a dose-dependent manner.

### Vaccination with MERS-1227 or MERS-1236 protects from viral shedding in alpacas

Alpacas have been established as a surrogate infection model for camels, the predominant natural intermediate animal host of MERS-CoV. Unlike camels, alpacas do not develop rhinorrhoea (nasal discharge) upon MERS-CoV infection, but they do shed infectious virus that is detectable in nasal turbinates^53,67^. We immunized alpacas (n = 5 per group) at two dose levels (20 µg or 200 µg) of MERS-1227 or MERS-1236 adjuvanted with 500 µg Alhydrogel using a prime-boost regimen (Figure 8A and Table S4). As a placebo control, an additional group of alpacas (n = 5) were dosed with 500 µg Alhydrogel at the same time points. Three weeks following the boost, (study day 41), all vaccinated alpacas had detectable pseudovirus neutralizing titers in their sera (Figure 8B). Furthermore, all alpacas vaccinated at both dose levels of MERS-1227 and the 200-µg dose of MERS-1236 had detectable neutralizing titers in a live-virus assay (Figure 8C).

**Figure 8.**
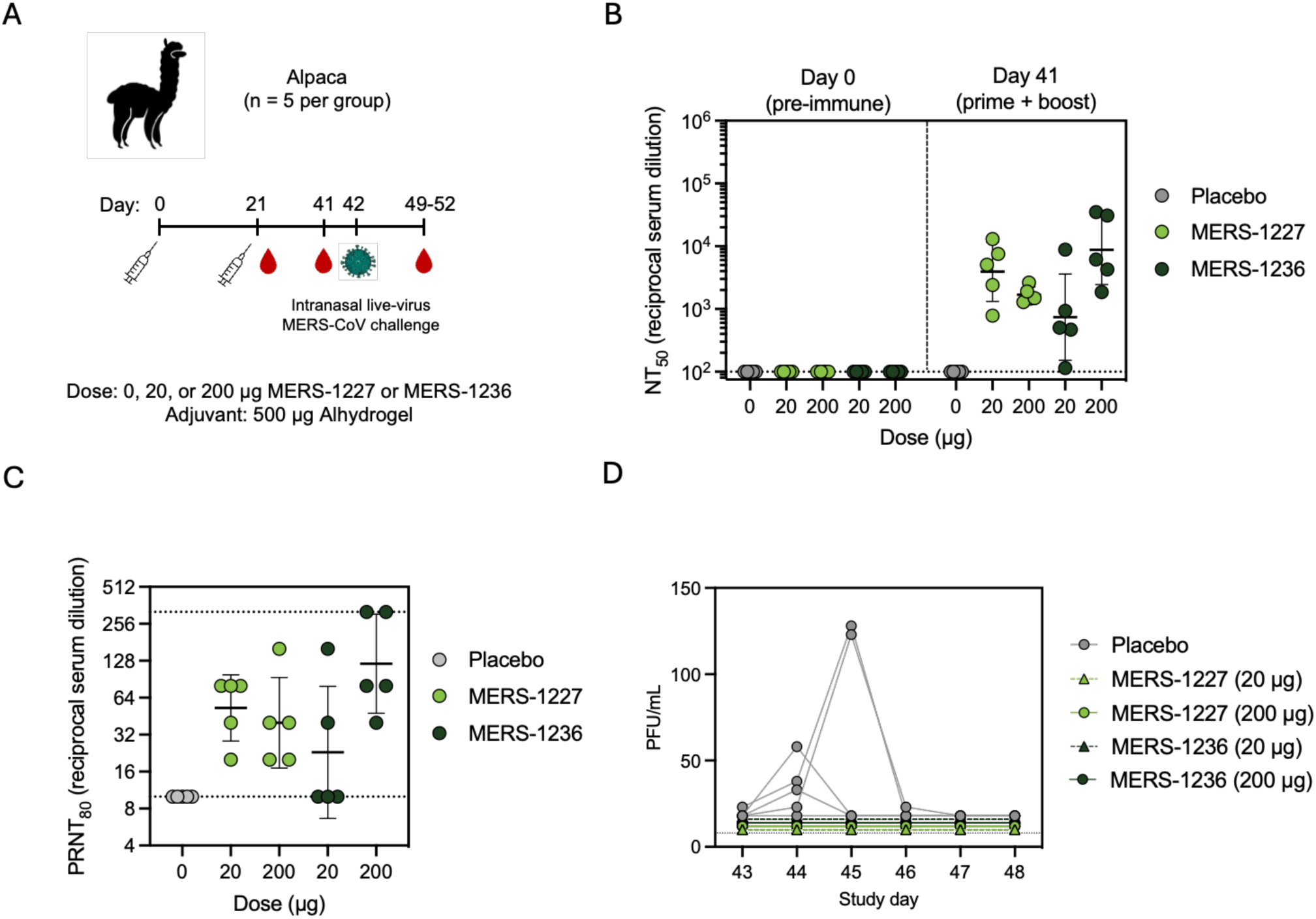
Alpacas immunized with Alhydrogel-adjuvanted MERS-1227 and MERS-1236 are fully protected from MERS-CoV infection. (A) Alpaca challenge study design and inoculation with EMC/2012 MERS-CoV. (B) Pseudovirus neutralizing titers from alpacas at day 0 (pre-immune) and 41 following two immunizations. Each circle represents the NT_5Ū_ value for an individual alpaca obtained by running pseudovirus neutralization analysis in quadruplicate. Lines represent the GMT and error bars represent the geometric SD. (C) Live-virus neutralization obtained from a plaque reduction neutralization assay using day 41 serum samples following two immunizations. Each circle represents the PRNT_80_ value for an individual alpaca. Lines represent GMT and error bars represent geometric SD. (D) Quantitation of live MERS-CoV virus in nasal swabs obtained from alpacas from days 1-5 and 7 post viral challenge via plaque assay. Each circle represents the mean number of plaque forming units observed in an individual alpaca from independent swabs of the left and right nare per mL inoculum.

On study day 42, alpacas were challenged by intranasal inoculation with live MERS-CoV virus (clade A, EMC/2012) in a BSL3 containment facility, as previously reported^67^. On days 1-5 and day 7 post-challenge, nasal swabs were collected from each nare, and live virus titers were determined using a plaque assay^67^. Detectable levels of MERS-CoV were observed via nasal swab for at least one time point following challenge in 4/5 alpacas that received the Alhydrogel placebo, while live virus was not detected in any (0/20) of the alpacas that received either dose level of Alhydrogel-adjuvanted MERS-1227 or MERS-1236 (Figure 8D). Taken together, these results demonstrate that immunization with Alhydrogel-adjuvanted MERS-1227 and MERS-1236 elicited protection from infection with live MERS-CoV. This provides strong evidence that these vaccines are protective and is an important demonstration that clinical development of this stabilized MERS-CoV spike ferritin vaccine could be utilized as a prophylactic countermeasure against MERS.

## Discussion

The severity of the COVID-19 pandemic has demonstrated the ongoing threat that emerging pathogens have on the global population and indicate the need for accessible prophylactic countermeasures. MERS-CoV, a betacoronavirus related to SARS-CoV-2 but with substantially greater morbidity and mortality, circulates in known animal reservoirs and causes sporadic outbreak clusters in humans with a case fatality rate (CFR) of over 30%^6^. Thus, development of a vaccine against MERS-CoV is a critical step towards pandemic preparedness in the event of a more widespread MERS-CoV outbreak. Here, we designed and evaluated two novel MERS-CoV vaccine candidates (MERS-1227 and MERS-1236), which display a stabilized form of the MERS-CoV spike on a ferritin nanoparticle. Our efforts primarily focused on a ferritin nanoparticle-based approach, a platform with a favorable safety, reactogenicity, and immunogenicity profile in humans^36,37^. While both MERS-1227 and MERS-1236 FNP proteins exhibited similar immunogenicity in mice, NHPs, and alpacas, MERS-1227 had more favorable structural and stability features as observed by TEM and DSF, and thus we focused our preclinical assessment on this candidate and used MERS-1236 as a comparator. The stability profile and immunogenicity of MERS-1227 FNP protein indicate that it is a strong candidate for progression into clinical development.

A number of key features differentiate the antigen design we describe here from previous nanoparticle-based MERS vaccine candidates. To our knowledge, we are the first to report an ectodomain truncation of the MERS-CoV spike; previously published work described either a full-length spike with the transmembrane domain included^68–70^ or a full-length ectodomain conjugated to a nanoparticle^71,72^. Given the biochemical differences we observe between MERS-1227 and MERS-1236 (Figure 2C-F), we hypothesize that inclusion of the additional residues of the ectodomain (residues 1237-1294) could lead to dramatic flexibility of the spike, which may impact neutralizing antibody responses and protein stability. Interestingly, the difference we observed in immunogenicity between MERS-CoV spike trimer and MERS-1227 and MERS-1236 (Figure 4C) did not appear in a recently published study from Chao et al. describing a MERS nanoparticle that contained residues 1237-1294^73^. This may suggest that the benefits of multimerization in our nanoparticle candidates are aided by the removal of this flexible domain and the stable presentation of the spikes on the surface of ferritin.

As an indication of the feasibility for scale-up and cGMP manufacturing of MERS-CoV spike ferritin nanoparticles, we showed that both candidates are expressed at high levels (∼ 2.7- 3.2 g/L titer) in CHO cells (Figure S1A) and can be purified via two orthogonal tag-less purification processes (Figure S1B). Furthermore, the unadjuvanted drug substances show a robust thermostability profile in which the molecules remain as folded, stable nanoparticles with retained antigenicity after 14 days at 37 °C (Figure 3). We also demonstrated nanoparticle stability following low pH treatment, which could be an important process development step required for viral clearance in cGMP manufacturing. These research studies show promise towards feasibility of developing a cGMP chemistry, manufacturing, and controls (CMC) process for these molecules.

The MERS-1227 and MERS-1236 FNPs are immunogenic in multiple animal models (BALB/c mice, NHPs, and alpacas). In BALB/c mice, we found that when adjuvanted only with Alhydrogel, MERS-1227 elicits neutralizing antibodies at doses as low as 0.016 µg (Figure S4). Furthermore, we tested multiple adjuvant combinations that markedly improved the immunogenicity of MERS-1227 (Figure 6A) demonstrating an opportunity to develop a clinical formulation to elicit the most robust and durable immune response. Finally, a key finding from our mouse immunogenicity studies was that encoding the MERS-1227 ferritin nanoparticle in mRNA elicits a comparable response to immunization with the Alhydrogel-formulated protein nanoparticle (Figure 6B). Given that protein subunit vaccines and mRNA vaccines have different advantages, it is valuable to observe that MERS-1227 elicits robust immunity after prime and boost injections in multiple platforms.

We evaluated the durability of the immune response to Alhydrogel-adjuvanted MERS- 1227 and MERS-1236 following a prime-boost regimen in NHPs with a three-month interval previously defined as an optimal boosting interval in other vaccine regimens^58^. These results revealed that a single dose of either MERS-1227 or MERS-1236 in NHPs elicits neutralizing antibodies that are quantifiable 3 weeks post vaccination and persist at a steady level until a second dose. The primary response is robustly boosted when a second dose is administered three months after priming (Figure 5B). Additionally, the neutralizing titers elicited following the boost are persistent and exhibit GMT values five months following the boost (day 253) that are ∼2-fold higher than the three-month post prime (day 93) timepoint. In addition to the durable response observed, serum from immunized NHPs inhibits in vitro infectivity from pseudotyped viruses displaying spike proteins from MERS-CoV clades A, B, C, and a distant pangolin merbecovirus (MjHKU4r-CoV-1)^57^ (Figure 5C). This suggests that in the event of crossover of clade C camel strains or potentially a more distant emergent merbecovirus into humans, this vaccine could provide protection. Furthermore, transgenic mice expressing human DPP4^66^ were protected from lethal MERS-CoV challenge following immunization with MERS-1227 in a dose-dependent manner and immunized animals exhibited reduced titers of live virus in the lung. Using an alpaca MERS-CoV challenge model^67^, we observed complete protection against viral shedding in animals vaccinated with either 20 µg or 200 µg of MERS-1227 or MERS-1236 adjuvanted with Alhydrogel. This provides strong evidence that these vaccines are effective at protecting against MERS-CoV infection.

Given the robust immune response we observed, including durable neutralizing antibody titers, prevention of infection in an established preclinical challenge model, and a favorable thermostability and manufacturing profile, we assess MERS-1227 as an optimal candidate for rapid clinical development. This would enable prophylactic vaccination for at-risk individuals, thus mitigating the risk of further MERS-CoV outbreaks, and if needed, for regional or global pandemic preparedness, readiness, and response.

## Methods

### Construct design and sequences

An England1 clade B sequence of MERS-CoV spike protein derived from AFY13307.1^23^ was truncated to reflect a similar C-terminal deletion as previously investigated for a SARS-CoV-2 spike^24,40^ and was stabilized using proline substitutions at residues 889, 1060, and 1061^20^. The S1/S2 furin cleavage site was mutated from the native RSVR to ASVG^20^. Given the lack of structural resolution at the C-terminal portion of the MERS-CoV spike ectodomain, two truncation positions were tested, one at position 1227 and one at 1236. An SGG linker was encoded following the truncation site prior to a modified *H. pylori* ferritin as previously described^24^. The terminal 3 residues in MERS-1227 encode a potential glycosylation site (NST). An N1225G mutation was made to remove this glycosylation site, as it was hypothesized that a glycan at the base of the trimer immediately preceding the ferritin component could impede nanoparticle formation.

### MERS-1227 and MERS-1236 stable CHO cell pool generation, fed-batch culture production, and purification

Stable CHO cell pools were generated as previously described^42^. Briefly, codon optimized coding sequences were synthesized and cloned into Leap-In™ transposon-based expression constructs. The plasmids were co-transfected with Leap-In™ transposase mRNA into miCHO-GS host cells (ATUM) and the stable pools were established by glutamine-free selection. For fed-batch productions, cells were seeded at 0.75 x 10^6^ per mL in shake flasks at either 25 mL or 100 mL scale grown at 37 °C with 5% CO2. Growth temperature was adjusted to 30 °C on day 4 and cells were supplemented throughout the fed-batch production using proprietary feed solutions. Cells were harvested on day 14 by spinning at 7,000 xg for 10-20 min. Supernatants were filtered using a 0.22-µm filter.

For research-scale purification, supernatants were diluted with 20 mM Tris pH 8.0, 1 M sodium chloride to a final concentration of 200 mM sodium chloride (not accounting for sodium chloride in the cell culture media). Diluted supernatants were then passed over a Cytiva 5 mL HiTrap Q anion exchange column equilibrated in 20 mM Tris pH 8.0 using an AKTA fast protein liquid chromatography (FPLC) system. MERS-CoV ferritin nanoparticles were collected from the column flow-through and then dialyzed overnight at 4 °C using cellulose ester 1 MDa molecular weight cutoff dialysis tubing (Spectrum) into 20 mM Tris, pH 7.5, 150 mM sodium chloride. Dialyzed material was concentrated to a final volume of ∼0.5-2 mL using an Amicon spin concentrator (100 kDa molecular weight cutoff). Concentrated samples were then filtered using a 0.22-µm filter and injected onto a Cytiva Superose 6 10/300 GL (PN 29091596) or Sepax 10 x 300 mm SRT SEC-1000 (particle size 5 µm, pore size 1000 Å, PN 215950-10030) size-exclusion chromatography column equilibrated in 20 mM Tris pH 7.5, 150 mM sodium chloride using an AKTA FPLC. Protein-containing fractions were assessed by SDS-PAGE analysis, and fractions containing homogenous protein were pooled. Fractions were supplemented with 5% sucrose as a cryoprotectant, frozen in liquid nitrogen, and stored at –80 °C until use. The formulation buffer used throughout the described studies was 20 mM Tris pH 7.5, 150 mM sodium chloride, 5% sucrose.

Alternatively, for bind and elute purifications (Figure S1B), MERS-1227 14-day fed-batch supernatant was treated with Pierce Universal Nuclease and then diluted 5-fold into 20 mM Tris pH 7.5. Diluted supernatant was injected onto a 1 mL Sartobind Q nano (Sartorius No.96IEXQ42DN-11--A) equilibrated in 20 mM Tris-Cl pH 7.5 at 5 mL per min. FNP was eluted using a step elution with 10 column volumes of 14, 30, and 100% 20 mM Tris-Cl pH 7.5, 1 M sodium chloride. FNP-containing fractions were pooled, brought to 1.6 M ammonium sulfate, and loaded onto a 1mL Cytiva HiTrap Hydrophobic Interaction Capto butyl column equilibrated in 1.8 M ammonium sulfate in 50 mM HEPES pH 7.5. The protein was eluted with a gradient of 50 mM HEPES pH 7.5.

### SDS-PAGE

Denaturing SDS-PAGE analysis was performed by diluting proteins with LDS sample buffer (containing 2-mercaptoethanol for reducing conditions) and heating samples for 5 min at 95 °C. Samples were then loaded on a 4-20% MiniPROTEAN TGX precast gel and run for 30 min at 230 V. Gels were stained using GelCode Blue staining reagent and imaged using a Thermo Fisher Scientific imager.

### Analytical size-exclusion chromatography multi-angle light scattering

Purified MERS-1227 and MERS-1236 were diluted to 0.125 - 0.5 mg/mL in 20 mM Tris, pH 7.5, 150 mM sodium chloride, 5% sucrose. For each run, 5-10 µg protein was loaded onto a 4.6 x 300 mm SRT SEC-1000 analytical column (particle size 5 µm, pore size 1000 Å, PN 215950-4630) pre-equilibrated in 20 mM Tris pH 7.5, 150 mM sodium chloride run at 0.35 mL/min on a Waters Arc Premier high-pressure liquid chromatography system. Multi-angle light scattering was measured using a Wyatt DAWN and refractive index was measured using a Wyatt Optilab. Molecular weight calculations were performed using Wyatt ASTRA software by utilizing the light scattering and refractive index signals from each sample.

### Sample preparation for LC-MS/MS peptide mapping

MERS-1227 and -1236 FNP proteins were proteolytically digested using S-Trap columns (Protifi, C02-micro-10) following the manufacturer protocol with minor modifications. All digestions were performed in triplicate. For each replicate, 25 µg protein in formulation buffer was added to an equal volume of 2x lysis buffer (5% SDS, 100 mM Tris, pH 8.5). Proteins were reduced in 5 mM dithiothreitol (DTT) for 30 min at 55 °C and alkylated in 20 mM iodoacetamide for 30 min in the dark at room temperature. Phosphoric acid was added to a final concentration of 2.5% (v/v). Acidified samples were diluted in 10 volumes of 100 mM Tris, pH 7.55 in 90% methanol/10% water and applied to S-Trap columns by centrifugation at 4,000 x g. The column was washed in the same buffer. 2.5 µg of trypsin (Promega, #V5113) diluted in 20 µL 50 mM ammonium bicarbonate, pH 8 was applied to the column and allowed to sit overnight at 37 °C. Peptides were eluted in 40 µL ammonium bicarbonate pH 8, followed by 40 µL 0.2% formic acid in water, and finally 40 µL 50% acetonitrile in water. Eluted peptides were dried on a SpeedVac and resuspended in 30 µL 0.1% formic acid in water prior to analysis.

### LC-MS/MS peptide mapping method

Peptides were analyzed by LC-MS/MS using a Vanquish Horizon LC directly interfaced to an Orbitrap Exploris 240 mass spectrometer (Thermo Fisher Scientific). 6 µL of resuspended peptides were separated by reversed-phase LC on a Accucore Vanquish C18+ (100 mm length x 2.1 mm ID, 1.5 µm particle size) at 0.3 mL/min using a linear gradient of 5-30% mobile phase B (0.1% formic acid in acetonitrile) over 25 min followed by a ramp to 90% mobile phase B over 3 min, where mobile phase A was 0.1% formic acid in water. Positive electrospray ionization was achieved with a spray voltage of 3500 V, a sheath gas flow rate of 40 arbitrary units, auxiliary gas heated to 250 °C at a flow rate of 15 arbitrary units, and a sweep gas flow rate of 1 arbitrary unit. The ion tube transfer temperature was held at 320 °C and the RF lens was set to 70. MS analysis was carried out in a Top 10 data dependent acquisition mode. Full MS scans were acquired at a resolution of 60,000 (FWHM @ m/z 200) with an automatic gain control target of “standard”, m/z range of 350–1800, and maximum ion accumulation time of “auto”. Precursor ions with charges 2–6 and a minimum intensity threshold of 1 x 10^4^ were selected with a quadrupole isolation window of 2.0 m/z for HCD fragmentation with a normalized collision energy (NCE) of 27%. MS/MS scans at a resolution of 15,000 (FWHM @ m/z 200) were acquired with an automatic gain control of “standard”, a maximum ion accumulation time of 150 ms, and a fixed first mass of 120 m/z. Precursors selected for fragmentation more than 2 times within a 15 second window were dynamically excluded from additional fragmentation for 10 seconds.

### LC-MS/MS peptide mapping data analysis

To identify post-translational modifications of MERS-1227 and -1236 ferritin nanoparticles, raw mass spectrometry data were analyzed using the GlycanFinder module of PEAKS Studio 11 (Bioinformatics Solutions, Inc). Custom FASTA databases were generated comprising the Chinese hamster (*Cricetulus griseus*) proteome (downloaded on November 27, 2023), common mass spectrometry contaminant proteins^74^, and either the MERS-1227 or -1236 FNP protein sequence. The raw MS data were searched against the database with the following modifications: fixed carbamidomethylation on Cys, variable oxidation of Met, and variable deamidation of Gln and Asn, with a maximum of two variable modifications per peptide. Glycans were identified using the default N-linked and O-linked databases within PEAKS GlycanFinder. Semi-specific tryptic peptides with a maximum of two missed cleavages were considered. The allowed mass tolerances were 10 ppm for precursor ions, 0.04 Da for peptide product ions, and 20 ppm for glycan diagnostic ions. Hits were filtered to a false discovery rate of 1% using the PEAKS decoy-fusion approach.

### Dynamic light scattering and differential scanning fluorimetry

Proteins were diluted to 0.125 - 0.5 mg/mL in formulation buffer (20 mM Tris pH 7.5, 150 mM sodium chloride, 5% sucrose) and loaded into glass capillaries (NanoTemper). Samples were then analyzed on a Nanotemper Prometheus Panta using DLS with a viscosity parameter of 1.149 mPa×s and a refractive index parameter of 1.341 to determine the cumulant radius of the particles and DSF to determine the melting temperature (Tm). Melting temperature runs were performed using a gradient of 1.0 °C per minute from 25 °C to 90 °C.

### Transmission electron microscopy

For TEM analysis, purified MERS-1227 and MERS-1236 were diluted using 20 mM Tris pH 7.5, 150 mM sodium chloride. A 3.5 μL drop of diluted sample suspension was applied to TEM grid overlaid with a 3-4 nm layer of amorphous C (CF200-CU-UL, Electron Microscopy Sciences). Prior to sample application, the carbon-coated grid was plasma-cleaned for 15 seconds via a lab-made device. The 3.5 µL of sample was allowed to incubate on the carbon surface for one minute. After blotting the sample away with filter paper, each grid was twice dipped quickly (∼1 second) into separate drops of de-ionized water followed by blotting with filter paper. This double washing and blotting was then repeated with two drops of 1% ammonium molybdate. For all steps, the next dipping was performed before the grid could completely dry. Finally, each grid was dipped onto a drop of 1% ammonium molybdate solution. After 15-20 seconds, the stain was blotted away with filter paper and the girds were allowed to air-dry. To collect hundreds of images rapidly, specimens were imaged on a ThermoFisher Titan Krios transmission electron microscope equipped with a Gatan Bioquantum K3 energy filter and direct electron detector. The microscope was operated at 300 kV and at liquid nitrogen temperatures, and the program SerialEM was used to collect images^75^. Images were collected as multiple frames of the same field-of-view.

After imaging, micrographs were processed to give two-dimensional class averages via the RELION software package^76^. Frames in the multi-framed images were aligned and summed via the MOTIONCOR2 algorithm^77^ implemented in RELION. Contract transfer function parameters were determined for each summed micrograph via CTFFIND4,^78^ via an interface in RELION. Several hundred particle images were selected manually and used to train the program Topaz (interfaced in RELION) to automatically pick thousands more particle images^79^. Extracted particle images were then aligned in two-dimensions to give 2D class averages via the general Bayesian approach of RELION^80^.

### Biolayer interferometry

Biolayer interferometry analysis of MERS-1227 and MERS-1236 was performed using an Octet R8 instrument with Protein A functionalized tips. Antibodies were procured from GenScript with human Fc domains and stored in 1X TBS pH 7.4. VH and VL sequences are found in Table S2.

Antibodies and MERS-CoV nanoparticles were diluted to 10 µg/mL in 1X Sartorius kinetics buffer (PBS + 0.1% BSA, 0.02% Tween-20, and a microbicide, Kathon) and pipetted into black-walled, black-bottom plates (200 µL per well). Protein A tips were dipped into antibody wells and subsequently dipped into nanoparticle wells to assess binding association for 120 sec. Dissociation was monitored for 120 sec by moving tips into a well containing buffer alone.

### pH and temperature stability treatment of MERS-CoV FNPs

To assess the pH stability of MERS-CoV FNP proteins, aliquots of the formulation buffer (20 mM Tris pH 7.5, 150 mM NaCl, 5% sucrose) were adjusted to pHs of 2, 3, 3.5, and 4 (+/- 0.1 pH unit) by dropwise addition of hydrochloric acid. Slide-A-Lyzer MINI Dialysis Devices (15 mL conical format, Thermo Fisher Scientific Cat #88401) were briefly rinsed in these buffer solutions. 350 µL of MERS-1227 or MERS-1236 FNP proteins at 0.25 mg/mL in formulation buffer were added to each dialysis device and dialyzed against 14.5 mL of the pH-adjusted formulation buffers (or pH 7.5 as a control). Proteins were dialyzed at room temperature for 1 hour with gentle agitation. After 1 hour incubation, 1 µL dialyzed protein was used to confirm the target pH of the solution using pH paper. At this time, dialysis solution was discarded and replaced with a fresh 14.5 mL pH-adjusted formulation buffer and dialysis was allowed to proceed for an additional 1.5 hours. Proteins were then dialyzed back to pH 7.5 by replacing dialysis solution with pH 7.5 formulation buffer followed by a 1 hour incubation at room temperature. pH 7.5 formulation buffer was then replaced with fresh pH 7.5 buffer and dialysis was allowed to proceed overnight at 4 °C. Proteins were then removed from dialysis devices, transferred to 2 mL polypropylene tubes, snap frozen using liquid nitrogen, and stored at -80 °C until analysis. For analysis, all samples were thawed at the same time and diluted to 0.125 mg/mL with formulation buffer and evaluated using analytical SEC, DLS, DSF, and BLI as described above.

To assess thermal stability of MERS-CoV FNP proteins, purified proteins were diluted to 0.5 mg/mL in 20 mM Tris pH 7.5, 150 mM NaCl, 5% sucrose and aliquoted in 2 mL polypropylene tubes. Samples were placed at either 4 °C, 22 °C (ambient temperature), or 37 °C. Control samples were snap-frozen immediately following dilution to 0.5 mg/mL and 90 °C heat-treated controls were heated to 90 °C for 30 min and then snap-frozen and stored at -80 °C until analysis. Aliquots of 4 °C, 22 °C, and 37 °C were taken following 7- and 14-day incubations and snap-frozen at each time-point until analysis. For analysis, all samples were thawed at the same time and assessed at 0.5 mg/mL using analytical SEC, DLS, DSF, and BLI as described above.

### Antigen production and purification

MERS-CoV GCN4 trimer, MERS-CoV RBD, and MERS-CoV NTD (amino acid sequences in Table S2) were produced in Expi293F and purified via His-tag purification. Expi293F cells were cultured in Expi293 media (Gibco) at 37 °C, 120 RPM, and 5% CO2. For transfection, cells were grown to a cell density of 3 x 10^6^ cells and transfected with plasmid DNA at 1 µg DNA per mL cell culture using ExpiFectamine293 transfection reagent (Gibco) according to manufacturer recommendations. Cells were harvested 4-6 days post transfection by centrifugation at 7000 x g followed by filtration using a 0.22-µm filter.

Filtered supernatant was diluted with 10X PBS to a final concentration of 2.5X PBS (pH 7.4) and loaded onto either a HisTrap excel (Cytiva) or a HiTrap TALON crude (Cytiva) column pre-equilibrated in 2.5X PBS pH 7.4. Proteins were eluted with 250 mM imidazole in 2.5X PBS pH 7.4. For Luminex, MERS-CoV GCN4 trimer was buffer exchanged into 20 mM Tris pH 7.5, 150 mM NaCl and biotinylated using a BirA biotinylation kit (Avidity). MERS-CoV RBD and NTD were purified on a Cytiva Superdex 200 Increase 100/300GL (PN 28990944) SEC column pre-equilibrated in 1X PBS pH 7.4. MERS-CoV GCN4 trimer was purified on a Cytiva S6 10/300 GL (PN 29091596) or a Cytiva Superdex 200 Increase 100/300GL (PN 28990944) SEC column pre-equilibrated in 20 mM Tris pH 7.5, 150 mM NaCl. Protein-containing fractions were pooled, snap-frozen, and stored at -80 °C until use.

### MERS-CoV FNP formulation for immunization studies

To formulate protein antigens with Alhydrogel adjuvant, the antigens were first diluted to the required concentration in formulation buffer (20 mM Tris pH 7.5, 150 mM NaCl, 5% sucrose). Alhydrogel adjuvant (Invivogen #vac-alu-10, 10 mg/mL) was resuspended by inverting 50 times, and then the required volume was added to the diluted antigen. The combined antigen-adjuvant mixture was then inverted 50 times, maintained at room temperature for 30-60 min to allow for antigen binding to Alhydrogel, and then inverted 50 additional times.

For the adjuvant screen (Figure 7A), MERS-1227 FNP was first formulated with Alhydrogel adjuvant as described above, except the FNP and Alhydrogel concentration were initially prepared to 2x the required final concentration for injection. For formulations without Alhydrogel, only the protein was included. Additional adjuvants were prepared as follows: ODN2395 (CpG, Invivogen Cat #trtl-2395) was dissolved to 1 mg/mL in sterile water, MPLA-SM (Invivogen Cat. #vac-mpla2) was dissolved to 1 mg/mL in sterile DMSO, and QuilA was dissolved to 1 mg/mL in sterile water. AddaVax (InvivoGen Cat t#vac-adx-10) was purchased as a 2x ready-to-use suspension. 3M- 052-AF was provided at 0.050 mg/mL by The Access to Advanced Health Institute (AAHI). For each formulation, the appropriate volume of adjuvant plus additional formulation buffer was added to bring the final concentration to that required for injection, and the formulated vaccines were inverted 50 additional times.

### Mouse immunization studies

Female BALB/c mice (age 7-8 weeks) were purchased from Charles River Laboratories and acclimated for at least one week in a contract vivarium (Fortis Life Sciences) prior to immunization. Antigens and adjuvants were formulated to deliver the indicated doses in 100 µL volume. Mice were injected intramuscularly with a 50 µL dose of antigen in each hind leg, to deliver 100 µL of vaccine formulation per immunization. To obtain blood via retroorbital bleeding, a capillary tube was inserted into the medial canthus. Blood was then transferred to an Eppendorf or BD Microtainer Blood Collection Tube (BD 365967), allowed to clot at room temperature for approximately 2 hours, and then spun down at 4000 rpm in a tabletop centrifuge. The serum layer was transferred to an Eppendorf tube and frozen at -80 °C. Prior to subsequent assays, serum was heat inactivated at 56 °C for 30 min. All mouse studies were conducted in accordance with IACUC-approved protocols.

### Luminex bead conjugation, total IgG binding and IgG subclass binding assays

To measure the total IgG response and IgG subclasses in mice and NHPs, we used Luminex assays adapted from a previously described method^81^. Biotinylated MERS-CoV GCN4 trimer (50 µg) was incubated with 2 x 10^6^ MagPlex®-Avidin Microspheres (Luminex #MA-A012-01) for 2 hours, then washed and resuspended in 1 mL Luminex wash buffer (0.1% BSA, 0.02% Tween- 20 in PBS pH 7.4). For other antigens (MERS-CoV RBD and MERS-CoV NTD), different MagPlex® Microspheres (Luminex #MC10026-01 to MC10029-01 and MC10034-01) were used. Activation involved 1 x 10^7^ microspheres with 50 mg/mL sulfo-NHS (Thermo Fisher Scientific #24510) and EDC (Thermo Fisher Scientific #22980) for 20 minutes, followed by adding 50 µg of antigen for a 2-hour incubation. Conjugated microspheres were resuspended in 1 mL Luminex wash buffer. The post-conjugation antigenicity of each antigen was validated by measuring binding to control mAbs: 3A3 (MERS-CoV GCN4 trimer S2)^49^, JC57-14 (RBD)^12^, and G2 (NTD)^47^.

Serum samples from mice and NHPs were diluted 50-fold in Luminex diluent (1% non-fat milk, 5% FBS, 0.05% Tween-20 in PBS pH 7.4). 25 µL of diluted serum was added to 25 µL of antigen-conjugated microsphere suspension (at least 1000 microspheres / antigen). After a 30- minute incubation, microspheres were washed three times with Luminex wash buffer. For total IgG detection, microspheres were incubated with 100 µL of goat anti-mouse IgG R-phycoerythrin (2 µg/mL, SouthernBiotech #1030-09) for 30 minutes. For mouse IgG subclasses, biotinylated secondary antibodies specific to IgG1 and IgG2a (4 µg/mL, SouthernBiotech #1070-08, #1080-08) were used, followed by 30 minutes with streptavidin-PE (5 µg/mL, Thermo Scientific #12- 4317-87). Results were read on a Luminex xMAP Intelliflex system.

### MERS-CoV spike-pseudotyped lentivirus production and viral neutralization assay

To assess elicitation of neutralizing antibodies by MERS-1227 and MERS-1236, we utilized a MERS-CoV spike-pseudotyped lentivirus neutralization assay. For virus production, HEK293T cells were cultured in cell-growth media (DMEM, 10% fetal bovine serum, 2 mM L-glutamine, 100 Units/mL penicillin, and 100 µg/mL streptomycin) and seeded at a density of 6 million cells in a 10-cm dish. One day following cell seeding, cells were transfected with the following plasmids:

3.4 µg MERS-CoV spike plasmid, 2.2 µg of each lentivirus helper plasmid (pHDM-Hgpm2, pHDM- tat1b, and pRC-CMV-rev1b)^51^, and 10 µg lentivirus packaging vector (pHAGE-CMV-Luc2-IRES-ZsGreen-W)^51^ using BioT transfection reagent. Lentivirus production plasmids were obtained through BEI Resources (NR-52948). For all pseudovirus neutralization panels with the exception of Figure 5C, the MERS-CoV spike plasmid encoded the spike protein (EMC/2012 strain) with the µ-phosphatase signal peptide (sequence in Table S2) and an 18 amino acid C-terminal truncation, which has previously been shown to improve viral titers^82^. For viruses used to generate the panel shown in Figure 5D, native sequences of all viral spike proteins were used, and an 18 amino acid C-terminal truncation was also included (sequences in Table S2). The media was removed and replaced with fresh media 18-24 hours post transfection. Viruses were harvested 72 hours post transfection by spinning at 300 xg for 5 min and filtering through a 0.45-µm PES filter. Viruses were frozen at -80 °C until use.

Infectivity assays with MERS-CoV spike-pseudotyped lentiviruses were performed in HeLa cells expressing human DPP4 (Cellecta). HeLa/DPP4 cells were plated in white-walled, white-bottom 96-well plates at 5,000 cells per well one day prior to or the day of infection. To quantify neutralizing antibody titers in serum from immunized mice, serum samples were diluted in cell-growth media. Serum dilutions were then mixed with MERS-CoV spike-pseudotyped lentivirus and virus/serum dilutions were incubated at 37 °C for 1 hr. Media was either aspirated off of HeLa/DPP4 cells and replaced with virus/serum dilutions or trypsinized HeLa/DPP4 cells were plated directly into virus/serum dilutions and left to incubate with cells for 72 hr at 37 °C. Infectivity in each well was quantified using BriteLite and luciferase signal was readout using a BioTek Synergy H1 microplate reader. Data was analyzed using GraphPad Prism. Control wells containing virus only and cells only were averaged, and serum dilution infectivity values were normalized to these averages set at 100% infectivity (virus only) and 0% infectivity (cells only). Normalized values were then fit with a 3-parameter non-linear regression (Y=Bottom + (Top- Bottom)/(1+(X/IC50)). For Figure 4B-C, statistical analysis was performed in GraphPad Prism using a two-way ANOVA with multiple comparisons.

### NHP immunization studies

Ten male cynomolgus macaques of Mauritian origin (*M. fascicularis*), 5 years of age, were used for immunization studies (Table S4). Animals were maintained at Alpha Genesis, Inc., and the study was approved by the Committee on the Care and Use of Laboratory Animal Resources (IACUC Approval: #23-7). Alhydrogel-adjuvanted MERS-1227 and -1236 were prepared to a final concentration of 0.1 mg/mL antigen and 1.5 mg/mL Alhydrogel adjuvant. NHPs were injected intramuscularly with 500 µL of vaccine formulation per immunization. To obtain blood, animals were sedated with ketamine HCL 10-20 mg/kg IM and blood was collected from the femoral vein using a 22g 1.5 inch needle, vacutainer sheath, and collection tube. Blood was allowed to clot at room temperature for approximately one hour and was then centrifuged at 2000 xg for 20 min. Serum was transferred to Eppendorf tubes, heat inactivated at 56 °C for 30 min, and frozen at - 80 °C.

### Design and procurement of MERS-1227 mRNA/LNP complexes for immunization

mRNA vaccines encoding MERS-CoV antigens (Figure 6B) were procured from GenScript. The MERS-1227 ferritin mRNA construct encoded the amino acid sequence described above for the ferritin nanoparticle protein. The MERS-1227 cell-anchored mRNA encoded for a MERS-CoV spike containing the England1 spike residues 1-1332 with a deletion of residues 1228-1295 to reflect the ectodomain deletion present in the MERS-1227 nanoparticle. This construct also included a GGS linker between residue 1227 and the transmembrane domain. The MERS-CoV FL cell-anchored mRNA contained the England1 residues 1-1332. Both of the cell-anchored mRNA constructs included the 3 proline mutations described above, a mutated furin cleavage site, and a cytoplasmic tail truncation to remove residues 1332-1353 which encode for a cytoplasmic retention motif^82,83^. Codons were optimized for mouse expression. RNAs were synthesized using N1-methyl-pseudouridine, capped with a 5’ Cap1, and stored in 1 mM sodium citrate pH 6.5 buffer until formulation. RNAs were formulated in SM102-containing LNPs at a concentration of 0.1-0.2 mg/mL and stored in PBS pH 7.4, 10% sucrose buffer at -80 °C until use.

### hDPP4 BALB/c mouse challenge study

DPP4 founder transgenic mice were obtained from Dr. Neeltje van Doremalen at Rocky Mountain Labs (NIH) and used to establish a colony at Colorado State University. Offspring were genotyped as described^66^. DPP4 heterozygotes and homozygotes were used in the challenge study and are indicated in Figure S4B.

Antigens and adjuvants were formulated to deliver the indicated doses in 100 µL volume. Mice were injected intramuscularly with a 50 µL dose of antigen in each hind leg, to deliver 100 µL of vaccine formulation per immunization. Blood was collected at study days 21 and 42 and sera were stored frozen until analysis. For challenge, mice were lightly anesthetized with ketamine-xylazine and challenged by intranasal instillation of 10,000 PFU of the EMC/2012 strain of MERS-CoV. Mice were weighed and scored for clinical signs of disease daily beginning immediately prior to challenge and extending to euthanasia. Half of the mice were euthanized 3 days post-infection and lungs collected for virus titration. The remainder of the mice were maintained up to day 10 post-infection to assess changes in body weight and survival.

### Alpaca challenge study

Twenty-five female alpacas, 5 to 9 years of age, were assigned randomly such that their ages were homogeneously distributed to 5 vaccine groups (Table S4) and identified by thermally sensitive microchip. Alpacas were immunized at study days 0 and 21 by intramuscular injection of Alhydrogel-adjuvanted nanoparticle or Alhydrogel only (placebo group). Two days prior to challenge on day 42, alpacas were moved into a large animal BSL3 facility. Blood was collected at the time of each immunization and on day 41, and sera were stored frozen until use.

Challenge was conducted as described previously^67^. The alpacas were sedated with xylazine and challenged by intranasal instillation of a total of 5.4 x 10^5^ plaque-forming units (back titrated dose) of the EMC/2012 strain of MERS-CoV, split into 1 mL per nare. Alpacas were evaluated clinically daily for 7 days post-challenge. A deeply inserted nasal swab from each side was collected daily on days 1 to 5 and 7 post-challenge. Swabs were broken off in tubes containing 1 mL of BA1 (Tris-buffered MEM containing 1% bovine albumin) supplemented with 5% fetal bovine serum and antibiotics (gentamycin, penicillin, and streptomycin), then stored frozen until assay. Alpacas were euthanized between 7 and 11 days post-challenge. Virus in nasal swab samples was titrated using a double overlay plaque assay on Vero cells^53^. Titers of neutralizing antibodies to MERS-CoV in sera were determined by plaque reduction neutralization^84^ and expressed as the reciprocal of the highest dilution resulting in 80% neutralization of virus.

## Acknowledgments

We acknowledge receipt of the plasmids used for lentivirus production from BEI Resources (NIH, NIAID), deposited by Dr. Jesse Bloom. We thank Dr. Neeltje van Doremalen for providing the hPPD4 transgenic founder mice. We would like to thank Peter S. Kim for input and advice on project design and influential comments and feedback on this manuscript. We would like to thank Daniel Stieh for review of this manuscript and helpful discussion on statistical analyses and data presentation. We would like to thank Leslie Goo, Adam Weiss, and Jeremy Huynh for helpful comments on this manuscript. We would like to thank Max Li for institutional support throughout the duration of this project.

## Author contributions

PAW designed MERS-CoV FNP constructs. AEP conducted fed-batch CHO productions. AEP and JO purified MERS-CoV FNPs. AEP, SP, and BAP designed and conducted stability studies on MERS-CoV FNPs. AEP and BJF expressed and purified antigens for Luminex studies and JLC designed and performed Luminex binding assays. BAP and JO performed LC-MS/MS experiments and BAP analyzed the peptide mapping data. DMB performed TEM imaging experiments and analysis. HC produced MERS-CoV spike-pseudotyped lentiviruses and designed and conducted lentivirus neutralization assays. AEP, VA, JEL, MSK, PAW, and BAP designed mouse and NHP immunization studies. AEP and BAP formulated antigens for mouse, NHP, and alpaca immunizations. VA designed and coordinated mRNA procurement for platform evaluation studies. CSD oversaw design of immunoassays used for FNP characterization. AW, AB, AH, and RB conducted live-virus neutralization assays and alpaca challenge study. AEP drafted the manuscript with input from BAP, MSK, and JEL. All authors reviewed the manuscript prior to submission.

---

**Figure S1.**
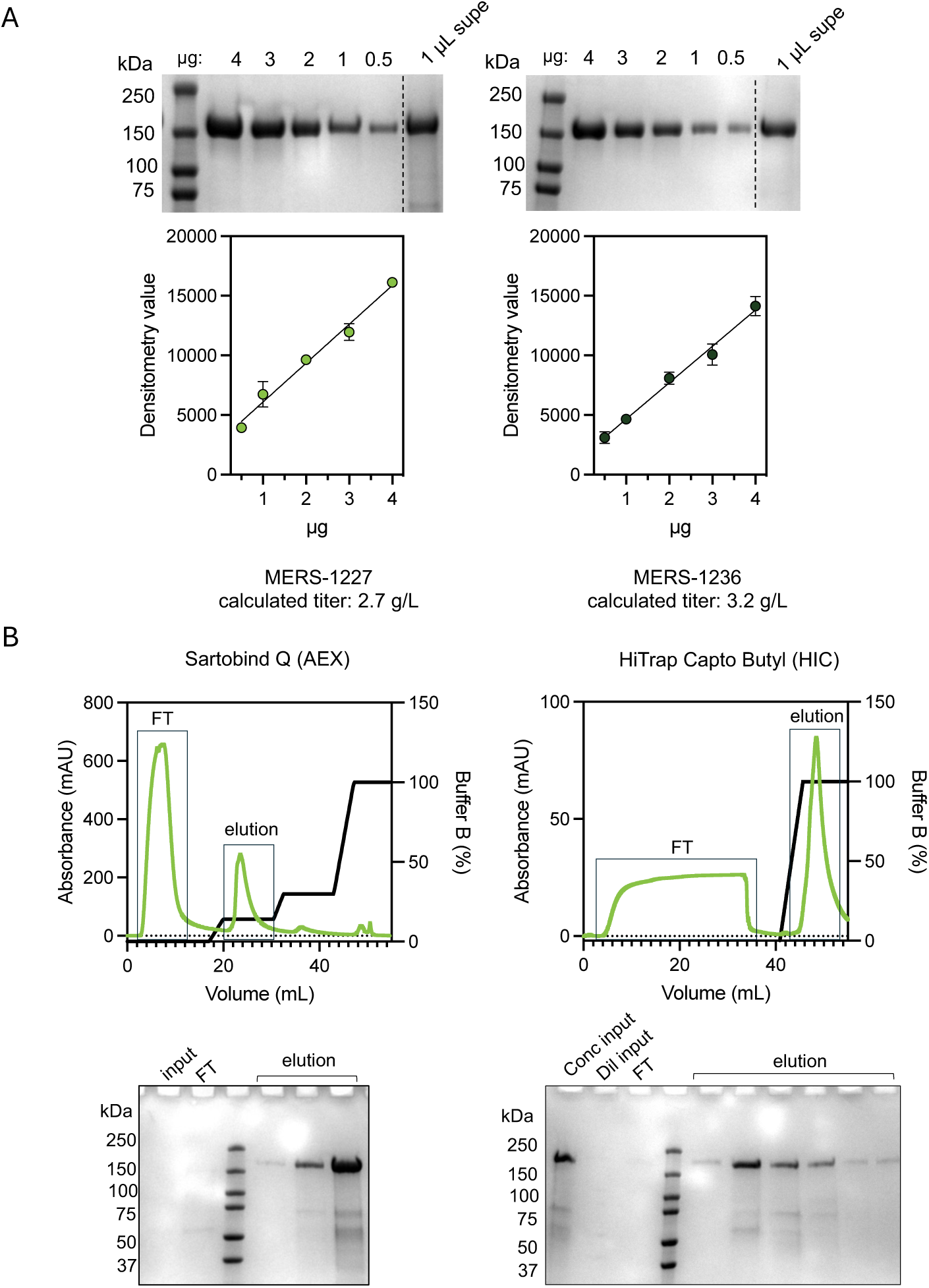
Quantitation of MERS FNPs in stable CHO pool supernatants and optimization of bind and elute chromatography purifications. (A) SDS-PAGE quantitation of MERS-1227 and MERS-1236 in 14- day fed-batch supernatants. Standard curves with 0.5-4 µg were generated using purified FNP proteins and quantified using densitometry. SDS-PAGE analysis was performed in duplicate; circles represent the mean and error bars represent the standard deviation. Densitometry was performed in ImageJ and quantitation was performed in GraphPad Prism. (B) Chromatograms obtained from AKTA purification on a Sartobind Q (left) and a HiTrap Capto Butyl (right) column. SDS-PAGE gels correspond to purification samples, with flow- through (FT) and elution fractions taken from the peaks boxed on the chromatograms above.

**Figure S2.**
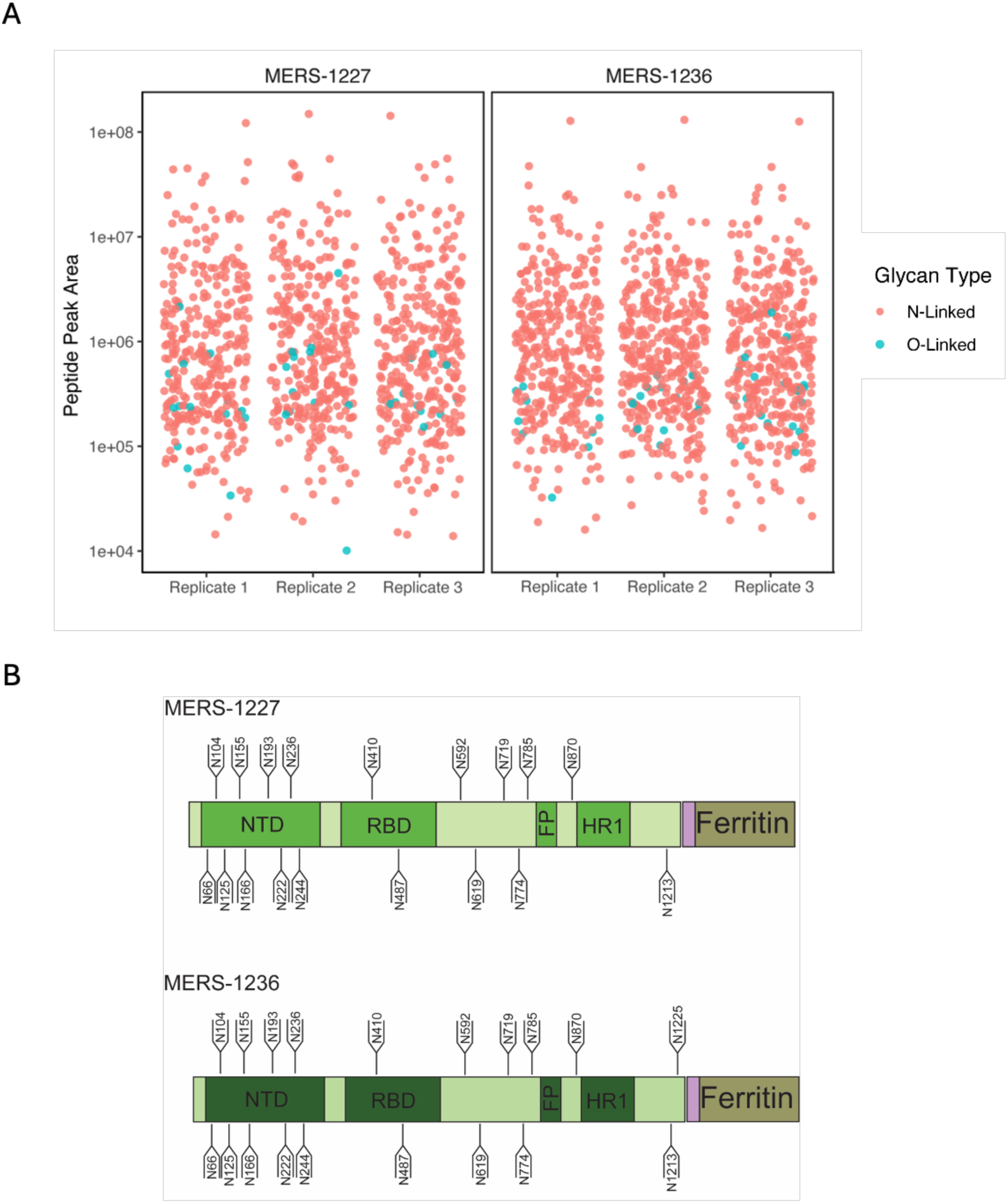
Characterization of MERS-1227 and MERS-1236 glycosylation using liquid chromatography­ mass spectrometry (LC-MS/MS). **(A)** Semi-quantitative assessment of N-linked vs. O-linked glycans on MERS-1227 and MERS-1236 FNPs. Individual peptides were assigned to have N-linked or O-linked glycans and their precursor peak areas were integrated using PEAKS GlycanFinder software. Each dot represents the area of an individual peptide. Samples were analyzed in triplicate. (B) Primary structure of MERS-1227 and MERS-1236 annotated with positions of detected N-glycosites. Full data on the N-glycans at each site, detected O-glycans, and summarized peak areas for each replicate are reported in Table S1.

**Figure S3.**
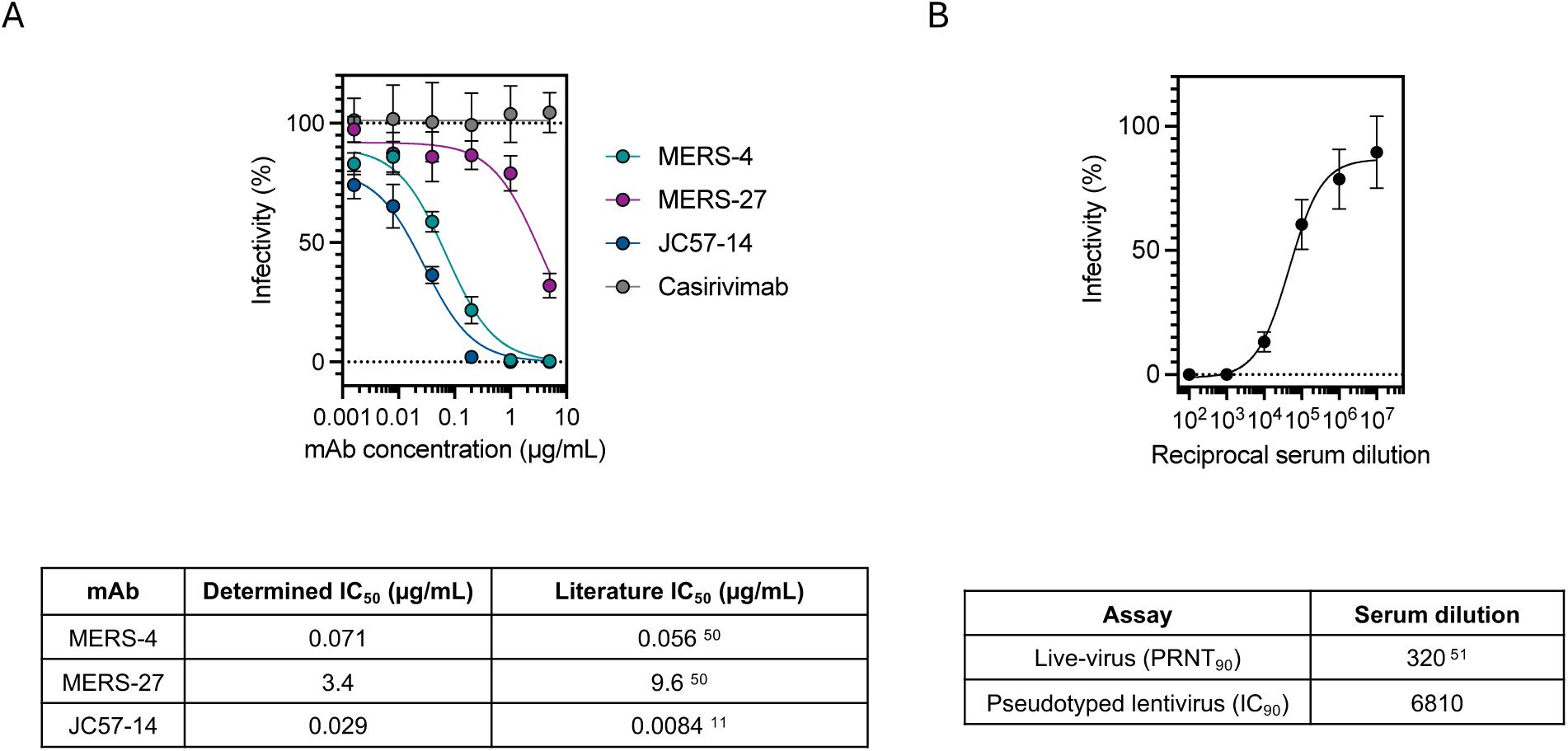
Validation of MERS-CoV spike-pseudotyped lentivirus neutralization assay using monoclonal antibodies and convalescent camel serum. (A) Pseudovirus neutralization assays run with serial dilutions of known MERS-CoV spike-targeting mAbs (list). A SARS-CoV-2 spike-targeting mAb (casirivimab) was included as a negative control. Samples were run in quadruplicate; circles represent the average % infectivity at each mAb concentration, and error bars represent the standard deviation. (B) Pseudovirus neutralization run with serum obtained from a camel 42 days following MERS-CoV challenge. Sample was run in n=10 replicates; circles represent the average % infectivity at each serum concentration, and error bars represent the standard deviation.

**Figure S4.**
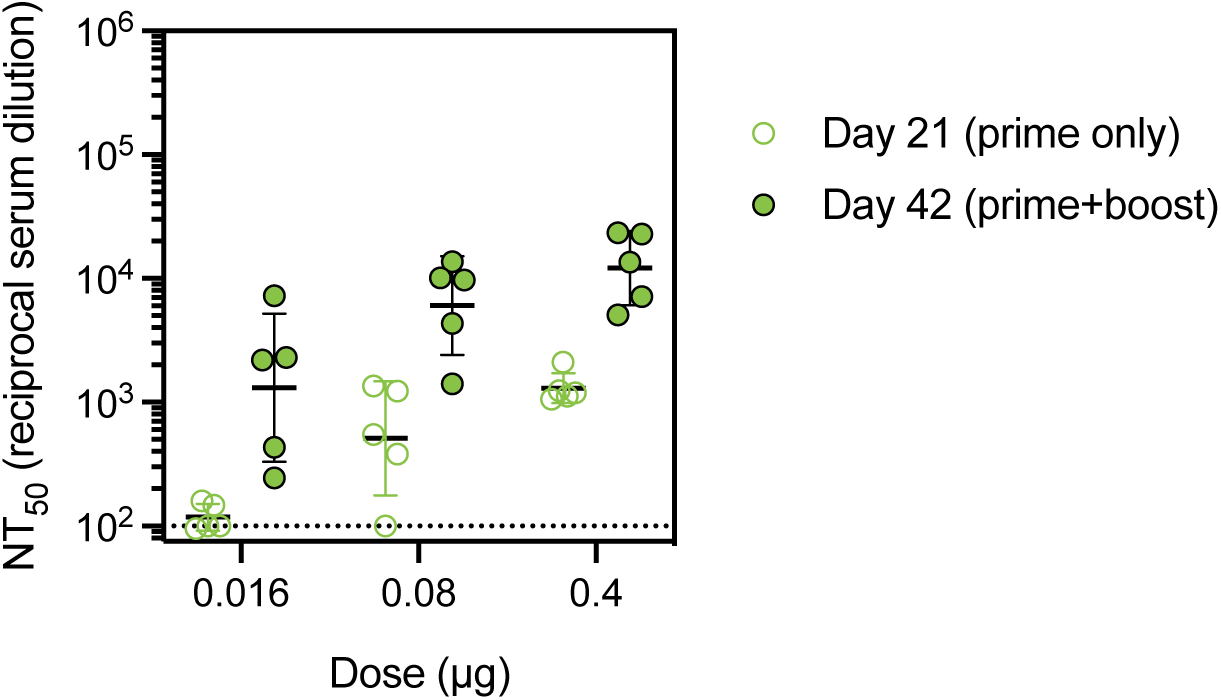
Dose de-escalation of MERS-1227 adjuvanted with Alhydrogel reveals quantifiable pseudovirus neutralizing titers following two doses as low as 0.016 µg FNP protein. Mice (n = 5) were immunized with 0.016, 0.08, or 0.4 µg MERS-1227 FNP protein adjuvanted with 150 µg Alhydrogel at days 0 and 21. Serum was collected at day 21 (prime only) and 42 (prime + boost) and assayed using pseudovirus neutralization. Each circle represents the NT_50_ for an individual mouse from quadruplicate measurements, bars represent the GMT, and error bars represent the geometric mean SD.

**Figure S5.**
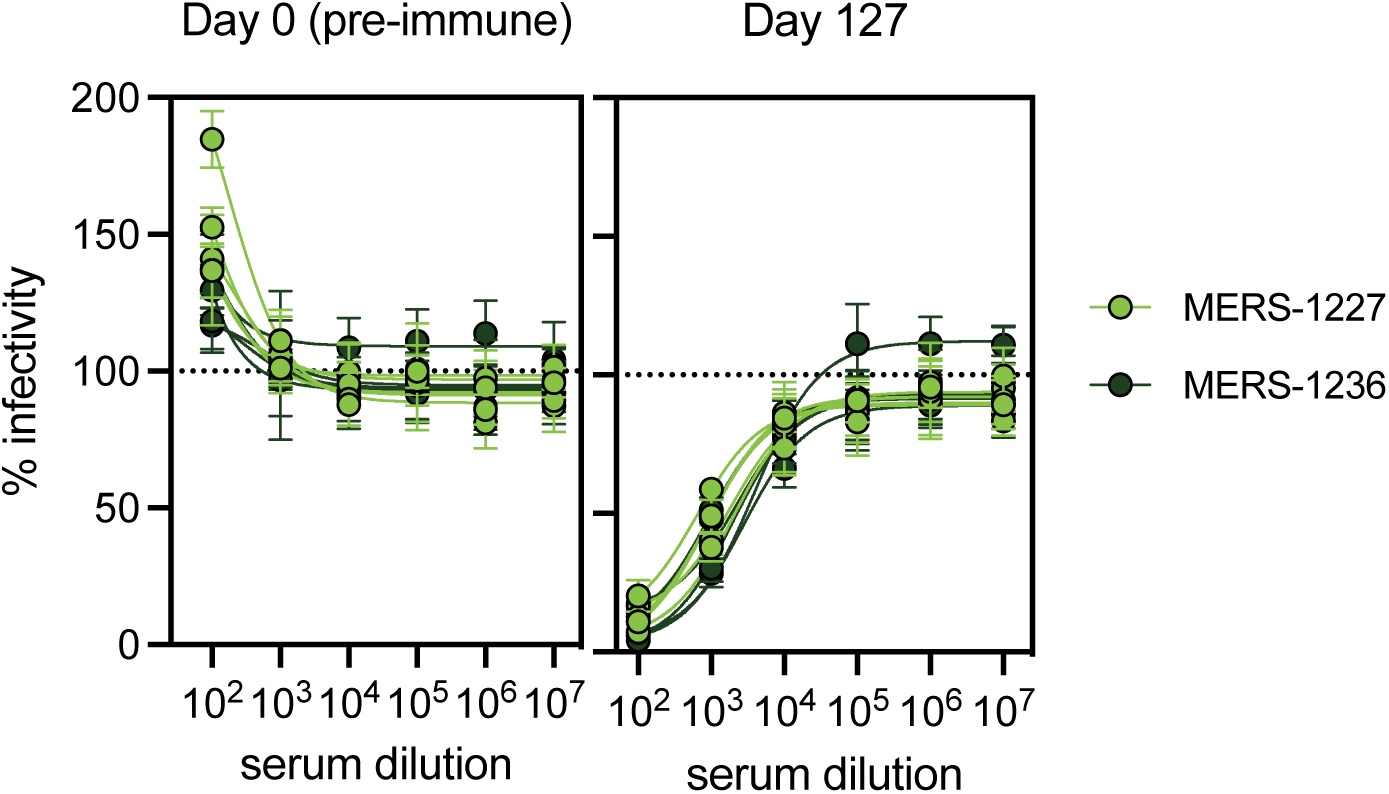
Pre-immune serum from NHPs does not neutralize lentivirus pseudotyped with MjHKU4r-CoV-1 spike. Full dilution curves from day 0 (pre-immune) and day 127 serum from NHPs shown in immunization study in Figure 5. Serum was assessed starting at a 1:100 dilution with 10-fold dilution steps. Each circle represents the mean % infectivity from quadruplicate measurements for a single NHP at each serum concentration and error bars represent standard deviation.

**Figure S6.**
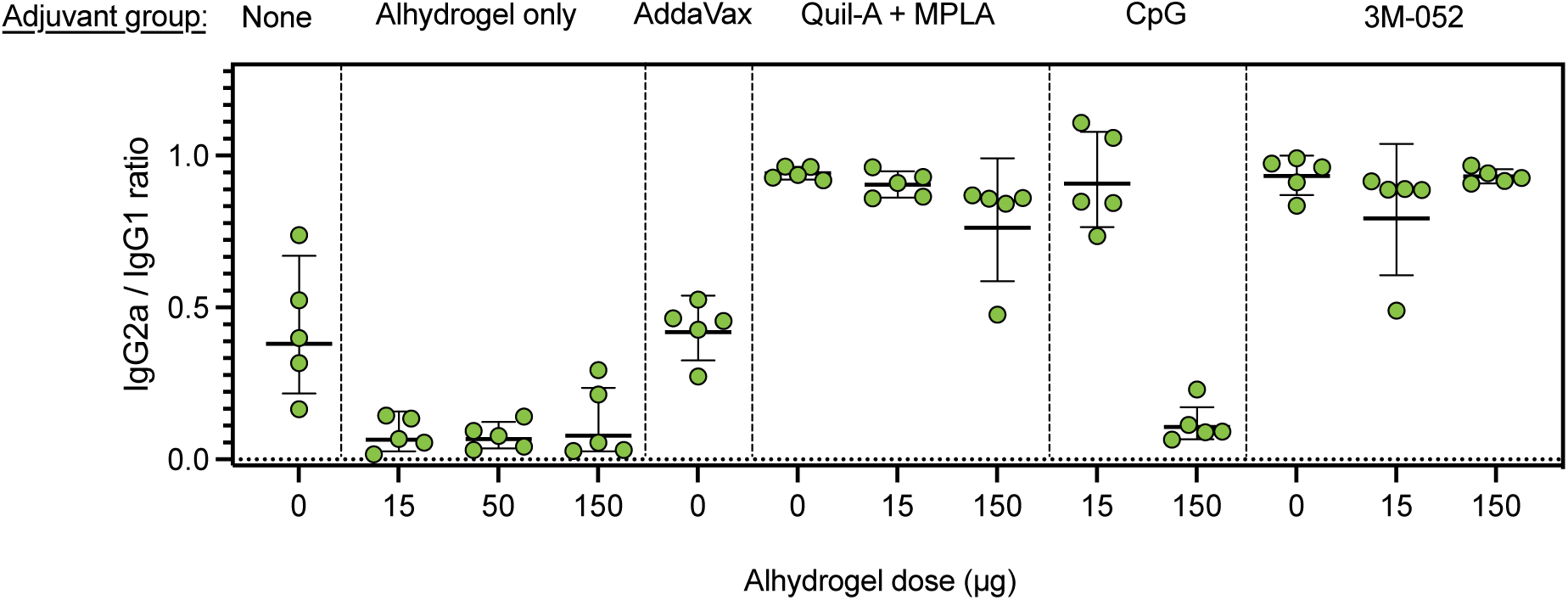
Ratio of IgG2a and IgG1 responses to MERS-CoV spike determined by Luminex reveal Th1/Th2 skewing effects of various adjuvants in BALB/c mice. Spike-specific IgG2a and IgG1 levels were determined using a Luminex binding assay for each mouse following two immunizations (day 42) with MERS-1227 formulated with adjuvant compositions indicated at the top. Binding was performed using biotinylated MERS-CoV trimer bound to streptavidin Luminex beads. The ratio was obtained by dividing the IgG2a value by the IgG1 value. The circles represent the ratio for each mouse, bars represent the geometric mean for each group, and error bars represent the geometric SD.

**Figure S7.**
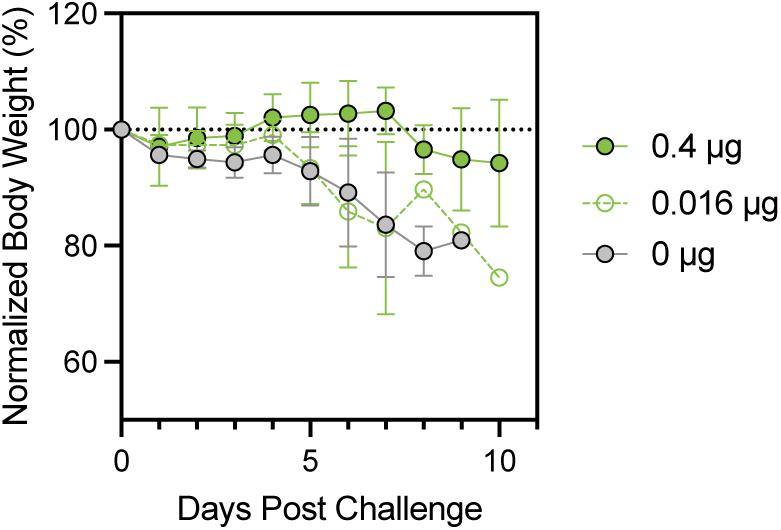
Weight loss following MERS-CoV challenge in hDPP4 mice immunized with MERS-1227. Mice (n = 3 or 4) were immunized with 0, 0.016, or 0.4 µg MERS-1227 FNP protein adjuvanted with 150 µg Alhydrogel at days 0 and 21 and challenged with EMC/2012 MERS-CoV at day 42. Mice were weighed daily post-challenge and weights are plotted as the averaged % body weight as compared to day 0 post-challenge. Each circle represents the mean weight per group at each timepoint and error bars represent SD. Mice that died were excluded from each time-point.

Table S1.**Glycan profiles of MERS-1227 and MERS-1236 FNP proteins.** Following tryptic digestions, peptides were analyzed by LC-MS/MS, and glycans were identified using PEAKS 11 GlycanFinder software. Identified glycan compositions at each glycosylated amino acid site are reported. Peak areas for each replicate reflect the extracted ion peak areas as determined by automatic integration by GlycanFinder. Entries with missing peak areas were not integrable, suggesting low abundance of these species.

Table S2.**Sequences of antibodies, recombinant proteins, and pseudovirus MERS-CoV spikes.**

**Table S3.** Pooled mouse serum from MERS-1227 and MERS-1236 immunizations neutralizes live MERS-CoV EMC/2012 via plaque reduction neutralization assay. Mouse serum taken at day 0 or 42 from the immunization shown in Figure 4 was pooled (40 µL per mouse) and evaluated using a live-virus PRNT assay. Serum dilutions corresponding to PRNT_90_, PRNT_80_, and PRNT_50_ are shown for each pooled serum sample.

| Serum pool | PRNT<sub>90</sub> | PRNT<sub>80</sub> | PRNT<sub>50</sub> |
| --- | --- | --- | --- |
| MERS-1227 Day 0 | <10 | <10 | 40 |
| MERS-1227 – Day 42 (0.4 µg dose) | >=320 | >=320 | >=320 |
| MERS-1227 – Day 42 (10 µg dose) | 160 | 160 | >=320 |
| MERS-1236 – Day 0 | <10 | <10 | <10 |
| MERS-1236 – Day 42 (0.4 µg dose) | >=320 | >=320 | >=320 |
| MERS-1236 – Day 42 (10 µg dose) | >=320 | >=320 | >=320 |

**Table S4.** Age and vaccine group for NHPs and alpacas included in immunization studies. (A) Ages and weights (kg) of NHPs immunized with MERS-1227 and MERS-1236 FNP proteins. (B) hDDP4 status (homozygous or heterozygous), MERS-1227 dose, and outcome for hDPP4 mice in MERS-CoV challenge study. (C) Ages and vaccine group designations for alpacas using in alpaca MERS-CoV challenge study.

| Identifier | Birthdate | Weight (kg) at time of first dose | Antigen |
| --- | --- | --- | --- |
| NV1702 | 15-Nov-17 | 5.22 | MERS-1227 |
| BC2312 | 26-Aug-18 | 4.54 | MERS-1227 |
| BC2483 | 30-Aug-18 | 4.5 | MERS-1236 |
| FR2573 | 6-Mar-18 | 6.14 | MERS-1236 |
| FR2864 | 2-Sep-18 | 4.56 | MERS-1236 |
| FR2913 | 8-Jul-18 | 6.02 | MERS-1227 |
| FR2923 | 9-Jul-18 | 4.36 | MERS-1227 |
| FR3151 | 27-Sep-18 | 4.4 | MERS-1227 |
| FR3188 | 29-Jul-18 | 4.16 | MERS-1236 |
| FR3267 | 3-Jan-18 | 4.37 | MERS-1236 |

B
| Identifier | Homozygous or heterozygous (HO/HZ) | Group | Dose (µg) | Day of euthanasia or death |
| --- | --- | --- | --- | --- |
| 34 | HZ | 1 | 0.16 | 3 (lung pathology) |
| 40 | HO | 1 | 0.16 | 3 (lung pathology) |
| 49 | HZ | 1 | 0.16 | 3 (lung pathology) |
| 56 | HZ | 1 | 0.16 | 3 (lung pathology) |
| 64 | HO | 1 | 0.16 | 7 |
| 74 | HZ | 1 | 0.16 | 7 |
| 80 | HZ | 1 | 0.16 | 10 |
| 87 | HZ | 1 | 0.16 | 7 |
| 35 | HO | 2 | 0.04 | 3 (lung pathology) |
| 41 | HZ | 2 | 0.04 | 3 (lung pathology) |
| 50 | HO | 2 | 0.04 | 3 (lung pathology) |
| 57 | HZ | 2 | 0.04 | 3 (lung pathology) |
| 68 | HZ | 2 | 0.04 | 10 |
| 75 | HZ | 2 | 0.04 | 10 |
| 81 | HZ | 2 | 0.04 | 8 (pregnant) |
| 88 | HO | 2 | 0.04 | 10 |
| 25 | HO | 5 | 0 | 3 (lung pathology) |
| 39 | HZ | 5 | 0 | 3 (lung pathology) |
| 48 | HO | 5 | 0 | 3 (lung pathology) |
| 55 | HZ | 5 | 0 | 3 (lung pathology) |
| 63 | HZ | 5 | 0 | 7 |
| 79 | HZ | 5 | 0 | 9 |
| 84 | HO | 5 | 0 | 8 |

| Identifier | Age (years) | Vaccine Group | Antigen and dose |
| --- | --- | --- | --- |
| 0892 | 6 | 1 | 20 µg MERS-1227 |
| 3529 | 7 | 1 | 20 µg MERS-1227 |
| 0890 | 8 | 1 | 20 µg MERS-1227 |
| 0897 | 7 | 1 | 20 µg MERS-1227 |
| 3526 | 9 | 1 | 20 µg MERS-1227 |
| 3523 | 5 | 2 | 200 µg MERS-1227 |
| 0895 | 7 | 2 | 200 µg MERS-1227 |
| 0879 | 9 | 2 | 200 µg MERS-1227 |
| 3528 | 7 | 2 | 200 µg MERS-1227 |
| 3527 | 9 | 2 | 200 µg MERS-1227 |
| 0893 | 6 | 3 | 20 µg MERS-1236 |
| 0896 | 7 | 3 | 20 µg MERS-1236 |
| 0898 | 9 | 3 | 20 µg MERS-1236 |
| 6532 | 7 | 3 | 20 µg MERS-1236 |
| 0891 | 7 | 3 | 20 µg MERS-1236 |
| 3525 | 4 | 4 | 200 µg MERS-1236 |
| 3524 | 8 | 4 | 200 µg MERS-1236 |
| 3522 | 7 | 4 | 200 µg MERS-1236 |
| 0889 | 7 | 4 | 200 µg MERS-1236 |
| 0894 | 9 | 4 | 200 µg MERS-1236 |
| 6530 | 6 | 5 | Placebo |
| 3521 | 7 | 5 | Placebo |
| 6533 | 7 | 5 | Placebo |
| 6531 | 7 | 5 | Placebo |
| 0880 | 9 | 5 | Placebo |

